# Human blood vessel organoids recapitulate key mechanisms of transition from vasculopathy to fibrosis in systemic sclerosis

**DOI:** 10.1101/2025.03.31.646441

**Authors:** Yanhua Xiao, Xuezhi Hong, Langxian Zhi, Yi-Nan Li, Martin Regensburger, Franz Marxreiter, Boris Görg, Sarah Koziel, Andrea-Hermina Györfi, Tim Filla, Peter-Martin Bruch, Philipp Tripal, James Adjaye, Sascha Dietrich, Jürgen Winkler, Jörg H.W. Distler, Alexandru-Emil Matei

**Affiliations:** Department of Rheumatology, University Hospital Düsseldorf, Medical Faculty of Heinrich Heine University, Düsseldorf, Germany; Hiller Research Center, University Hospital Düsseldorf, Medical Faculty of Heinrich Heine University, Düsseldorf, Germany; Department of Internal Medicine 3, Rheumatology and Clinical Immunology, Friedrich-Alexander-University Erlangen-Nürnberg (FAU) and University Hospital Erlangen, Erlangen, Germany; Deutsches Zentrum Immuntherapie (DZI), University Hospital Erlangen, Kussmaulallee 4, 91054, Erlangen, Germany; Spatial and Functional Screening Core Facility, Medical Faculty of Heinrich Heine University, Düsseldorf, Germany; Department of Stem Cell Biology, Friedrich-Alexander-Universität (FAU) Erlangen-Nürnberg, Erlangen, Germany; Center for Rare Diseases Erlangen (ZSEER), University Hospital Erlangen, FAU Erlangen-Nürnberg, Erlangen, Germany; Department of Molecular Neurology, FAU Erlangen-Nürnberg, Erlangen, Germany; Clinic for Gastroenterology, Hepatology, and Infectiology, Heinrich-Heine University, Düsseldorf, Germany; Department of Haematology, Oncology and Clinical Immunology, University Hospital Düsseldorf, Germany; Center for Integrated Oncology Aachen-Bonn-Cologne-Düsseldorf, Aachen Bonn Cologne, Germany; Molecular Medicine Partnership Unit, Heidelberg, Germany; Düsseldorf School of Oncology, Germany; Fraunhofer Institute for Translational Medicine and Pharmacology ITMP, and Fraunhofer Cluster of Excellence for Immune Mediated Diseases CIMD, Frankfurt am Main, Germany; Department of Hematology, Oncology and Rheumatology, University Hospital Heidelberg, Germany; Optical Imaging Competence Centre, Friedrich-Alexander-Universität Erlangen-Nürnberg, Erlangen, 91054, Germany; Institute for Stem Cell Research and Regenerative Medicine, University Hospital Düsseldorf, Moorenstrasse 5, D-40225 Duesseldorf, Germany; Zayed Centre for Research into Rare Diseases in Children (ZCR), University College London (UCL)-EGA Institute for Women’s Health, 20 Guilford Street, London WC1N 1DZ, UK

**Keywords:** Systemic sclerosis, microvasculopathy, blood vessel organoids, induced pluripotent stem cells, EndMT, Co-detection by indexing, multi-omics

## Abstract

Systemic sclerosis (SSc) is an autoimmune disease that transitions from vasculopathy as an initiating pathogenic event to tissue fibrosis. The mechanisms of these transitions remain, however, poorly understood, mainly because complex multicellular human models of SSc vasculopathy are lacking.

Here we characterized blood vessel organoids (BVOs) as a novel model system of vasculopathy in SSc. We demonstrate that exposure of SSc-BVOs to SSc serum triggers changes on epigenetic, mRNA and protein levels and recapitulates key pathogenic features of SSc vasculopathy, with shifts from angiogenic endothelial cell subsets to those undergoing endothelial-to-mesenchymal transition, loss of endothelial cells-pericytes interactions and profound angiogenic defects. The genetic predisposition of SSc donors and serum IgGs are required for the deleterious effects of SSc serum. We further validate SSc-BVOs as a human model system to evaluate candidate therapies targeting SSc microvasculopathy and use this system to provide evidence that γ-secretase inhibition is a potential therapeutic approach.

## Introduction

Connective tissue diseases, also known as collagen vascular diseases, are autoimmune disorders within the rheumatic spectrum, characterized by vasculopathy, inflammation, and tissue fibrosis as shared pathogenic features (1, 2). Systemic sclerosis (SSc) is the connective tissue disease associated with the highest case-specific mortality of all autoimmune rheumatic diseases (3). SSc is characterized by widespread vasculopathy, autoimmunity and fibrosis (4, 5). Microvasculopathy is the first detectable manifestation of SSc (6), and is thought to be the initiating event that triggers autoimmunity and fibrosis, but also plays a crucial role in disease progression (7). SSc is thus a prototypical connective tissue disease that transitions from microvasculopathy to inflammation and tissue fibrosis.

Endothelial cell damage triggers a cascade of pathologic events including aggregation of platelets at the denuded basal membrane, recruitment of leukocytes and subsequent activation of fibroblasts. Microvascular changes are visualized in the clinical routine by nailfold capillaroscopy. In early stages of SSc, characteristic findings include capillary dilatation and distorted capillary architecture, whereas in advanced disease, these microvascular changes progress to loss of capillaries and small arteries, leading to avascular areas (8, 9).

The vascular pathogenesis of SSc remains incompletely understood. Endothelial cell dysfunction is thought to result from the interplay of inflammatory features such as autoantibodies and cytokines with a subsequent imbalance between pro-angiogenic and anti-angiogenic factors in genetically predisposed individuals (10–15). Several susceptibility genes have been implicated in endothelial damage and vascular pathology in SSc, including polymorphisms in genes related to angiogenesis or oxygen sensing (16–19). However, the relative contribution of the individual drivers and their interplay remains elusive.

Our limited understanding of the vascular pathophysiology of SSc is at least in part a consequence of the paucity of preclinical models. Microvascular endothelial cells are more challenging to isolate from fibrotic tissues of SSc patients than many other cell types such as fibroblasts. Moreover, most mouse models of SSc do not recapitulate the microvascular features of SSc and the few mouse models that display certain vascular features of SSc are difficult to breed and are available only at few centers (20, 21). Multicellular human in-vitro-models of the vascular changes in SSc with sufficient complexity have not been reported so far.

In this study, we aimed to assess whether blood vessel organoids (BVOs) derived from pluripotent stem cells (iPSCs) of SSc patients and healthy individuals exposed to serum from patients with and without severe microvasculopathy can replicate key features of SSc vasculopathy. These BVOs are composed of endothelial cells and pericytes which self-assemble in three-dimensional vessel structures. We conducted a comprehensive multi-omics analysis to investigate how SSc derived iPSCs and SSc serum induce an SSc-like vascular phenotype. Finally, we explored the potential of BVOs as a novel platform for testing vasoactive drugs for the treatment of SSc.

## Results

### Generation of SSc- and healthy-BVOs and comparison of their vessel structure

We first generated BVOs from SSc-iPSCs and control-iPSCs (SSc-BVO and healthy BVO, respectively) following a previously described differentiation method (22, 23). This involved reproducing vessel development in embryonal life, in several steps: 1) formation of aggregates from iPSC, 2) mesoderm induction, 3) vascular lineage induction with formation of vessel aggregates and 4) collection of single vascular networks and further vascular differentiation to form BVOs (Fig. 1A). Confocal microscopy confirmed the successful formation of vascular networks in both groups, consisting of CD31 endothelial cells, PDGFRβ and/or SM22+ pericytes, and a surrounding COLIV basement membrane (Fig. 1B-C, Supplementary Video 1 and 2, Supplementary Fig. 1A). Three-dimensional (3D) image reconstitution revealed intricate vascular structures in both healthy and SSc BVOs, where endothelial cells formed tubular networks in close association with pericytes and the basement membrane (Fig. 1C, Supplementary Video 2).

**Figure 1.**
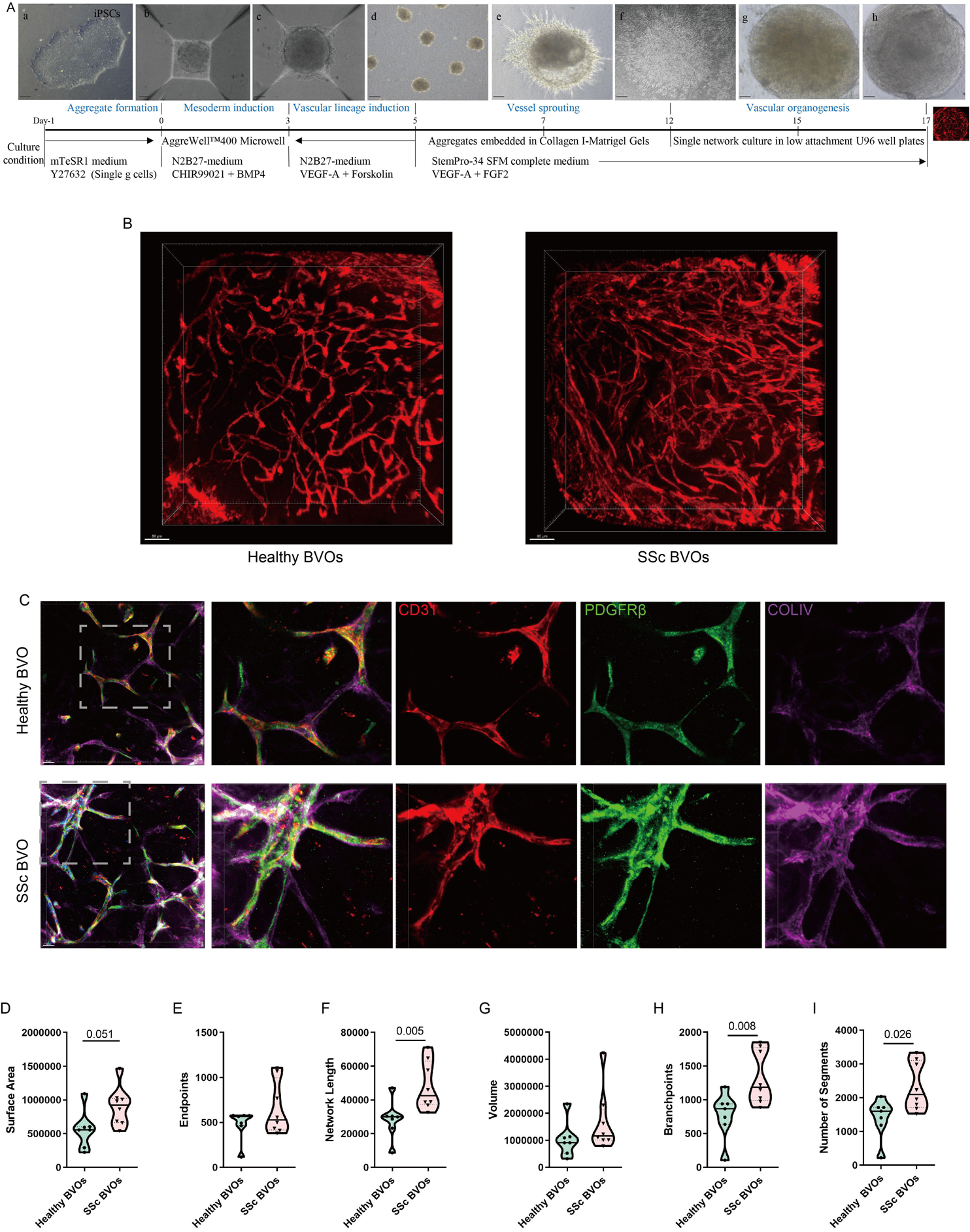
Generation and characterization of BVOs derived from healthy and SSc iPSCs. (**A**) Schematic representation of the protocol used for the differentiation of human pluripotent stem cells into blood vessel organoids. Representative phase contrast images at different timepoints during differentiation are included. Scale bars: 100 μm. (**B**) 3D rendering of confocal images of healthy and SSc-derived BVOs. CD31 immunostaining (red) identifies endothelial cells forming tubular vascular structures. Scale bars: 80 μm. (**C**) Confocal immunofluorescence of vascular networks in healthy and SSc BVOs, showing endothelial cells (CD31, red), pericytes (PDGFRβ, green), and basement membrane (COLIV, purple) (maximum intensity projection). Scale bars: 30 μm. (**D-I**) Quantitative comparison of vascular parameters between healthy and SSc BVOs, including surface area (D), endpoints (E), network length (F), volume (G), branchpoints (H), and number of segments (I), analyzed using VesselVio and ImageJ, and shown as violin plots. Each condition included four different clones with 1–2 organoids each. Statisctically significant *p*-values (Mann-Whitney U test) are included. SSc: systemic sclerosis; BVO: blood vessel organoids; 3D: three-dimensional.

We next analyzed vessel morphology using two tools: 1) AngioTool (24), which relies on two-dimensional maximum intensity projection representations of the 3D images, and 2) VesselVio (25), which uses directly 3D images, to quantify vessel density, length, volume and degree of branching, which are parameters commonly evaluated in nailfold capillaroscopy in clinical practice. SSc BVOs had a higher vessel density and total vessel length, thus indicating an upregulation of angiogenesis (Supplementary Fig. 1B-G). However, the SSc BVOs also had a higher number of branching points and of vessel segments, as well as a higher average volume and a higher percentage of vessels with very high volume, indicating a lower degree of maturation of the newly formed vessels (Fig. 1D-I). This reflects early changes in SSc vasculopathy, characterized by aberrant angiogenesis with formation of highly branched immature vessels and presence of giant capillaries (10, 26).

### Exposure to serum from SSc patients with severe vasculopathy induces angiogenic defects in SSc-BVOs

While aberrant angiogenesis with formation of immature vessels is typical for early stages of SSc, later disease stages are marked by angiogenic defects with extensive loss of small vessels (27). We reasoned that circulating factors in SSc patients might be required to reproduce this phenotype in SSc BVO. To test this hypothesis, we exposed SSc and healthy BVOs to serum from SSc patients suffering from active ischemic digital ulcers as clinical manifestation of severe ongoing microangiopathy at the time of serum prelevation (denoted as SSc_aDU serum) and serum from healthy donors (denoted as healthy serum) for seven days. Exposure to serum from SSc patients without active digital ulcers (denoted as SSc_noDU serum) was included to control for the effects of other disease-associated mediators (e.g. proinflammatory and profibrotic) on angiogenesis.

SSc_aDU serum induced profound angiogenic defects in SSc BVOs, as evidenced by a reduction in vessel density, vessel length, number of junctions, branching index, surface area and volume occupied by vessels (Fig. 2A-G, Supplementary Fig. 2A-F). In contrast, SSc_noDU serum caused only mild vascular changes (Fig. 2A-G, Supplementary Fig. 2A-F), suggesting that SSc BVOs can recapitulate the severity of vasculopathy present in the serum of donors. Interestingly, neither SSc_aDU nor SSc_noDU serum induced major vasculopathic features in healthy BVOs (Fig. 2A-G, Supplementary Fig. 2A-F). This suggests that SSc BVOs are particularly vulnerable to SSc_aDU serum-induced vasculopathy. These findings highlight that the combination of genetic predisposition and serum-mediated effects are required to fully manifest SSc-associated microvascular changes in BVOs.

**Figure 2.**
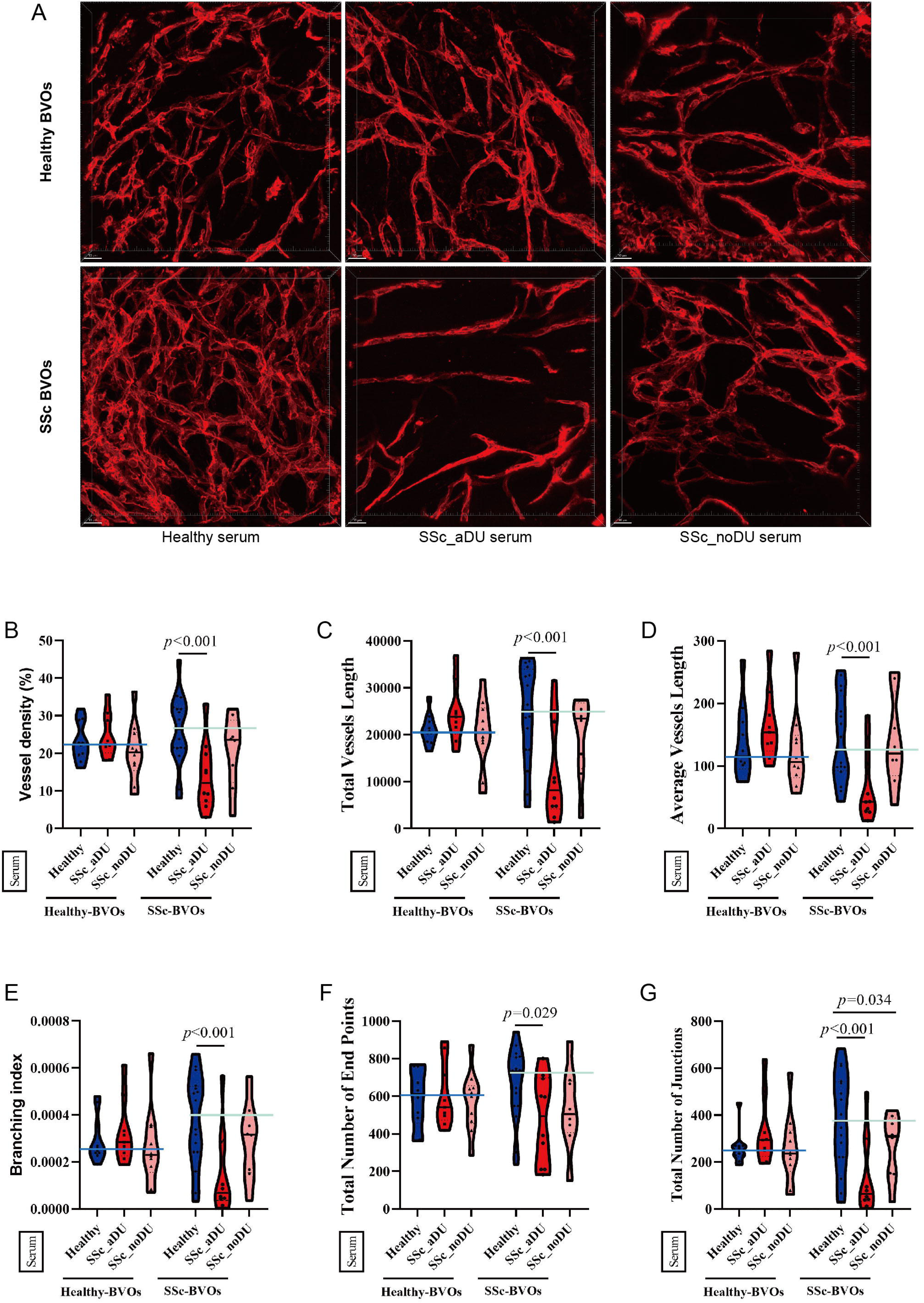
Serum from patients with clinically manifest microangiopathy induces angiogenic defects in SSc BVOs. (**A**) Representative 3D rendering of confocal images of healthy and SSc-derived BVOs treated with healthy serum, SSc_aDU serum, or SSc_noDU serum for seven days. Endothelial cells forming vessel structures were identified by CD31 marker expression (red). Scale bars: 30 μm. (**B-G**) Quantitative analysis of vessel parameters using AngioTool, including vessel density (B), total vessel length (C), average vessel length (D), branching index (E), total number of endpoints (F) and total number of junctions (G), shown as violin plots. Each serum treatment condition included four different clones with 2–4 organoids each. *P*-values are included from LMMs with serum treatment as a fixed effect and random intercepts fitted for each cell clone identity. SSc: systemic sclerosis; BVO: blood vessel organoids; 3D: three-dimensional; SSc_aDU serum: serum from SSc patients with active digital ulcers; SSc_noDU serum: serum from patients without active digital ulcers; LMM: linear mixed-effects model.

### Multi-omics-based characterization of SSc serum-induced vasculopathy in SSc BVOs

To characterize the mechanisms that lead to the profound angiogenic defects in SSc BVO exposed to SSc_aDU serum, we next performed a multi-omics-based evaluation of SSc BVOs exposed to SSc_aDU and healthy serum at the epigenomic level by ATACseq, at the transcriptomic level by RNAseq and at a multiplexed protein level by Co-detection by indexing (CODEX).

Exposure to SSc_aDU serum induced both changes in chromatin accessibility, as well as in transcriptomes of SSc BVOs treated with SSc_aDU serum, with 612 genes with differentially accessible regions (DAR) (104 with gain and 498 with loss of DAR), as well as 94 differentially expressed genes (DEGs) (24 upregulated and 70 downregulated) (Fig. 3A, B).

**Figure 3.**
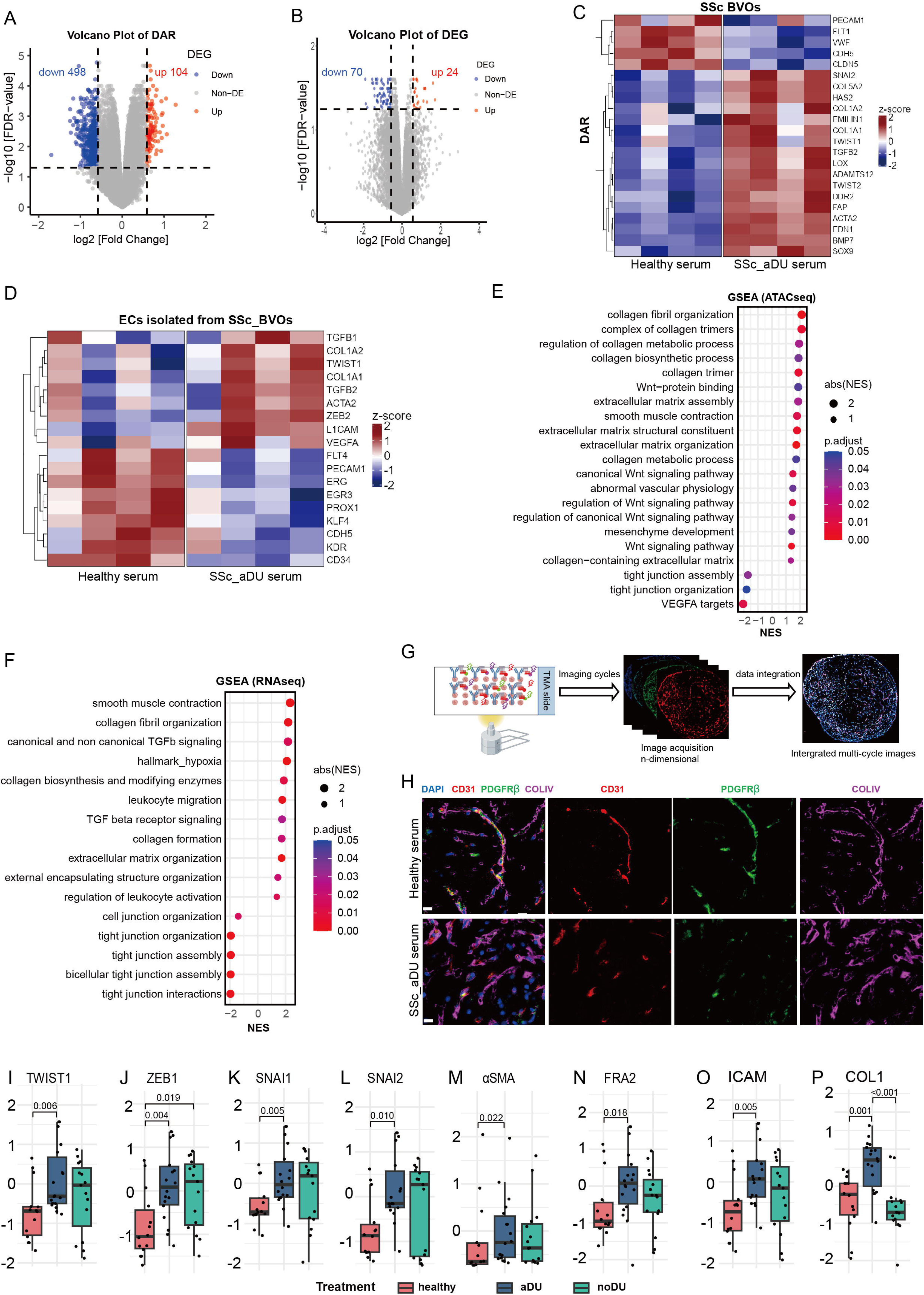
Multi-omics-based characterization of SSc serum-induced vasculopathy in SSc BVOs. (**A**) Volcano plot of ATACseq-based DARs in SSc BVOs exposed to SSc_aDU serum treatment compared to SSc BVOs exposed to healthy serum. Red and blue dots indicate genes with significantly upregulated and downregulated accessible regions, respectively ((|log2 fold-change| > 0.585; FDR < 0.05). (**B**) Volcano plot of RNAseq-based DEGs between SSc BVOs treated with SSc_aDU serum and healthy serum. (**C-D**) Heatmap of EndMT-associated markers identified from ATAC-seq data of whole BVOs (C), and from RNA-seq data of ECs isolated from SSc-BVOs (D), treated with SSc_aDU serum or healthy serum. (**E**) Bubble plot of GSEA based on ATACseq counts showing significantly enriched pathways in SSc BVOs treated with SSc_aDU serum compared to healthy serum. (**F**) Bubble plot of GSEA based on RNAseq counts between SSc_aDU serum and healthy serum-treated SSc BVOs. (**G**) Schematic illustration of multi-cycle imaging acquisition and data integration using CODEX technology. (**H**) Representative CODEX images of vascular structures of SSc BVOs treated with healthy serum or SSc_aDU serum. Markers include CD31 (red, endothelial marker), PDGFRβ (green, pericyte marker), and COLIV (magenta, basement membrane marker). Nuclei are stained with DAPI (blue). Scale bar, 20 µm. (**I-P**) CODEX-based protein expression levels of TWIST1 (I), ZEB1 (J), SNAI1 (K), SNAI2 (L), αSMA (M), FRA2 (N), ICAM (O), and COL1 (P) in SSc BVOs treated with healthy serum, SSc_aDU serum or SSc_noDU serum (n≥14 from four iPCS clones), shown as box plots (median ± IQR). Statistical significance was determined by the Kruskal-Wallis test followed by the Dunn multiple comparisons test. SSc: systemic sclerosis; BVO: blood vessel organoids; DAR: differentially accessible region; DEG: differentially expressed gene; SSc_aDU serum: serum from SSc patients with active digital ulcers; ATACseq: assay for transposase-accessible chromatin sequencing; RNAseq: RNA sequencing; EndMT: endothelial-to-mesenchymal transition; GSEA: Gene Set Enrichment Analysis; CODEX: co-detection by indexing; IQR: interquartile range.

Notably, several pro-fibrotic genes, including *ACTA2*, *TWIST1*, *SNAI2*, *TGFB2* or *COL1A2* gained DARs and were upregulated at the mRNA level in endothelial cells purified from SSc BVOs, while several endothelial cell marker genes, such as *CLDN5*, *PECAM1*, *CDH5*, and *VWF*, lost DAR and were downregulated at the mRNA level in endothelial cells purified from SSc BVOs, suggesting a shift toward a pro-fibrotic and EndMT-like phenotype in SSc BVOs treated with SSc_aDU serum (Fig. 3C, D).

Pathway enrichment analysis revealed consistent de-enrichment of pathways related to tight junction, endothelial integrity, and vascular function, thus indicating a compromised endothelial phenotype, with enrichment of pathways related to collagen production, extracellular matrix organization, contraction, TGFβ and WNT signaling and leukocyte migration and activation in SSc BVOs exposed to SSc_aDU serum both at the epigenomic and at the transcriptomic level (Fig. 3E, F, Supplementary Fig. 3A, B). Furthermore, transcription factor (TF) activity inference from the RNAseq data provided evidence for decreased activity of TFs related to an endothelial cell phenotype, such as GATA2, ETS1, FOXO4, NFATC2, ELK1 and ELK4 in endothelial cells purified from SSc BVOs, and increased activity of pro-fibrotic TFs including JUN, RELA, SMAD3, SNAI1 in SSc BVOs exposed to SSc_aDU serum (Supplementary Fig. 3C, D).

CODEX analysis confirmed that SSc_aDU serum induces angiogenic defects in SSc BVOs (Fig. 3G, H) and demonstrated upregulation of endothelial-to-mesenchymal transition (EndMT) markers (Twist1, ZEB1, SNAI1, SNAI2), myofibroblast markers (αSMA), pro-fibrotic transcription factors (FRA2), and increased expression of collagen type I at protein level (Fig. 3I-P). Furthermore, SSc_aDU serum induces upregulation of ICAM1, an endothelial cell activation marker, in SSc BVOs (Fig. 3O).

Of note, SSc_noDU serum treatment had milder pro-EndMT effects in SSc BVOs than SSc_aDU serum, whereas neither SSc_aDU nor SSc_noDU serum induced EndMT in healthy BVOs (Fig. 3I-P, Supplementary Fig. 3E).

Our multi-omics approach thus highlights that exposure to SSc serum from patients with severe microvascular disease induces changes on the epigenetic, mRNA and protein levels to induce a vascular phenotype in SSc BVOs that reproduces key pathogenic features of SSc vasculopathy, including EndMT.

### SSc BVOs recapitulate endothelial cell and pericyte populations observed in SSc skin

We next aimed to leverage the single-cell resolution of CODEX and evaluate changes in subpopulations of endothelial cells and pericytes and their function in SSc BVO upon treatment with SSc serum (Fig. 4A). We analyzed 70,660 cells, including 50,619 from SSc BVOs and 20,041 from healthy BVOs, each group exposed to either healthy, SSc_aDU and SSc_noDU serum.

**Figure 4.**
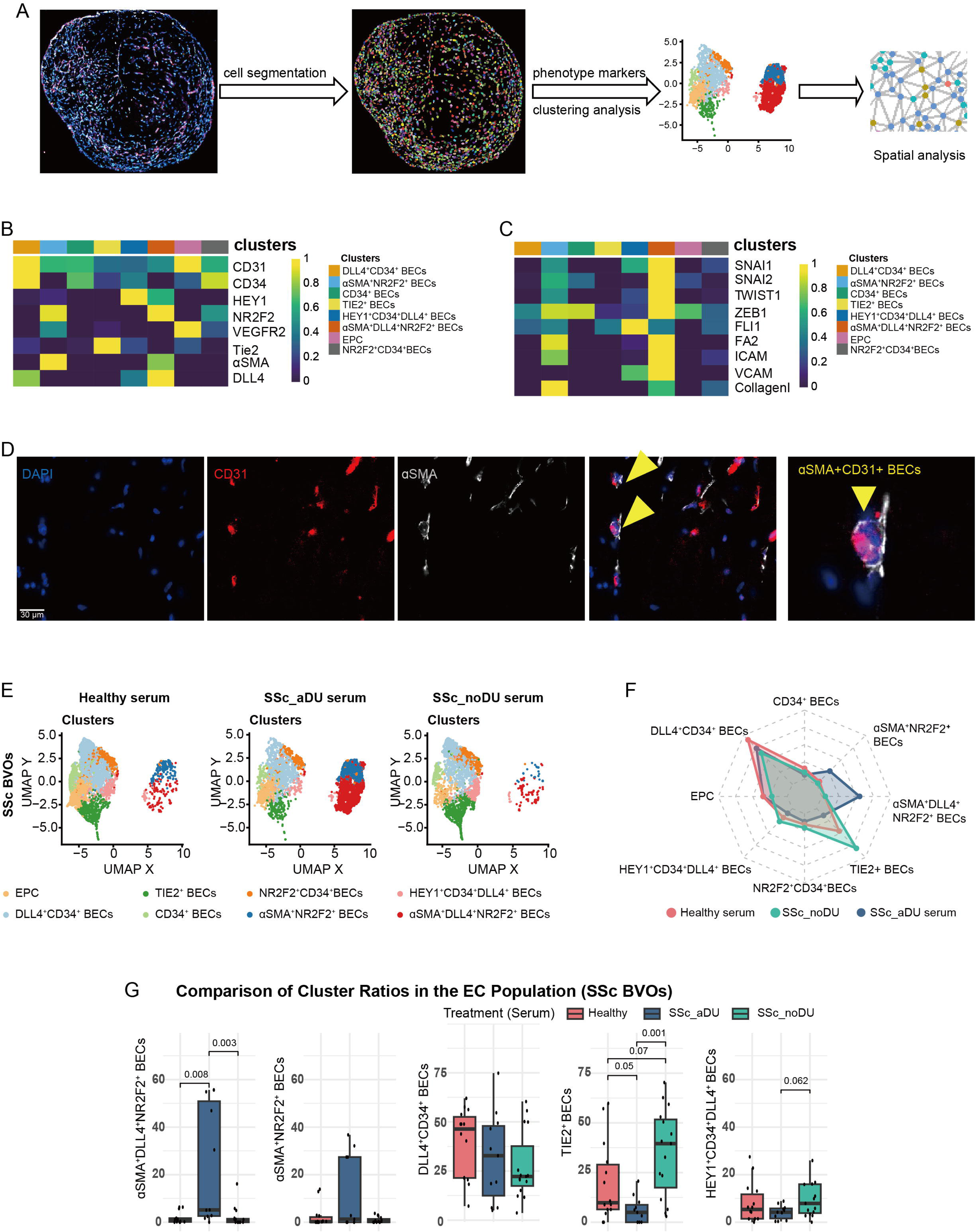
CODEX-based characterization of endothelial cell subpopulations in SSc BVOs and their shifts upon treatment with SSc serum. (**A**) Schematic workflow of CODEX single cell data processing and analysis. (**B, C**) Heatmap showing the relative expression levels of endothelial phenotype markers (B) and EndMT markers (C) across BEC subpopulations identified by CODEX analysis. (**D**) Representative CODEX images of CD31 (red), αSMA (grey) and DAPI (blue). Arrowheads indicate αSMA^+^CD31^+^ BECs. Scale bars = 30 µm. (**E**) UMAP plots showing the BEC subpopulations in SSc BVOs treated with healthy serum, SSc_aDU serum, or SSc_noDU serum. (**F**) Radar chart showing the changes in frequencies among BEC subpopulations (from all BECs) upon exposure to healthy serum, SSc_aDU serum and SSc_noDU serum in SSc BVOs. (**G**) Changes in frequencies of BEC subpopulations in SSc BVOs exposed to healthy serum, SSc_aDU serum or SSc_noDU serum (n≥11 from four iPCS clones) shown as box plots (median ± IQR). Statistically significant *p*-values (Kruskal-Wallis test followed by Dunn’s multiple comparisons test) are included. SSc: systemic sclerosis; BVO: blood vessel organoids; CODEX: co-detection by indexing; SSc_aDU serum: serum from SSc patients with active digital ulcers; SSc_noDU serum: serum from SSc patients without active digital ulcers; EndMT: endothelial-to-mesenchymal transition; BEC: blood endothelial cell; UMAP: Uniform Manifold Approximation and Projection; IQR: interquartile range.

We performed metaclustering of all cells to identify pericytes (PEs), expressing high levels of PDGFRβ, RGS5, CD90, SM22, and αSMA, but no CD31; lymphatic endothelial cells (LECs), expressing CD31 and high levels of LYVE1, VEGFR2 and VEGFR3, and blood endothelial cells (BECs), with strong expression of CD31, but with no or low levels of the pericyte-and LEC-specific markers (Supplementary Fig. 4A, B).

We further aimed to evaluate whether SSc BVOs recapitulate BEC subpopulations identified in SSc skin by single cell analyses in previous studies (28–30). We subclustered the BECs and identified eight distinct BEC subpopulations: 1) DLL4^+^CD34^+^ BECs, with an endothelial tip cell phenotype; 2) CD34^+^ BECs; 3) HEY1^+^DLL4^+^CD34^+^ BECs, demonstrating arterial BEC commitment; 4) NR2F2^+^CD34^+^ BECs, with venous BEC commitment; 5) TIE2^+^ BECs, with an endothelial stalk cell phenotype; 6) endothelial progenitor cells (EPCs), expressing VEGFR2 and CD34, 7) αSMA^+^DLL4^+^NR2F2^+^ BECs and 8) aSMA NR2F2 BECs, expressing high levels of αSMA as a marker of EndMT (Fig. 4B, D) (31). Thus, BVOs reproduce key endothelial cell populations previously described in SSc and healthy vasculature, including arterial BECs, venous BECs, stalk and tip cell BECs, endothelial progenitor cells, αSMA^+^ BECs and LECs (28–30).

The two αSMA^+^ EC populations expressed high levels of other EndMT-associated markers (SNAI1, SNAI2, ZEB1, TWIST1) as well as high levels of intracellular collagen type 1, and are thus bona fide endothelial cell subsets undergoing EndMT (Fig. 4C). Furthermore, they expressed high levels of the endothelial cell activation markers ICAM and VCAM, high levels of FRA2 and low levels of FLI1 (Fig. 4C), an expression pattern reminiscent of a dysfunctional endothelial cell phenotype previously described in SSc (32).

We further subclustered the pericytes to evaluate whether SSc BVOs recover pericyte populations observed in SSc skin (28). We identified four distinct populations: 1) CD90^hi^ PEs; 2) aSMA^hi^SM22^hi^ PEs; 3) PDGFRβ^hi^ PEs and 4) ADAM12^hi^ PEs (Supplementary Fig. 5A, B).

BVOs thus recapitulate previously described pericyte populations in SSc skin, including CD90^hi^, PDGFRβ^hi^ and ADAM12^hi^ PEs, as well as a population of aSMA^hi^SM22^hi^ PEs that demonstrate evidence of pericyte-to-myofibroblast transition (28).

### SSc BVOs exposed to SSc serum recapitulate shifts in frequencies of endothelial cell and pericyte populations observed in SSc skin

We further evaluated whether SSc_aDU serum induces shifts in frequencies of BEC and PE subsets in SSc BVOs. We observed remarkable shifts in BEC subpopulations following SSc_aDU serum treatment in SSc BVOs, with increases in frequencies of the two aSMA populations that are undergoing EndMT and decreases in endothelial cell populations involved in angiogenesis (DLL4 CD34 BECs, Tie2 BECs, and HEY1 CD34 DLL4 BECs) (Fig. 4E-G). Of note, enrichment of an aSMA subpopulation of endothelial cells undergoing EndMT was previously described in SSc skin (28).

We further observed an increase in the frequency of the αSMA^hi^SM22^hi^ and ADAM12^hi^ PEs and a decrease in the frequency of PDGFRβ^hi^ and CD90^hi^ PEs upon exposure of SSc BVOs to SSc_aDU serum (Supplementary Fig. 5B, C). Enrichment of ADAM12^hi^ and αSMA^hi^ pericytes, as well as loss of CD90^hi^ pericytes was previously described in SSc skin (28). Taken together, these results demonstrate that exposure of SSc BVO to SSc_aDU serum reproduces shifts in endothelial cell and pericyte subsets observed in SSc skin.

Of note, the shifts in EC and PE subpopulations in response to SSc_aDU serum are specific for SSc BVOs and are not observed in healthy BVOs (Supplementary Fig.4D, Supplementary Fig.5D).

### Altered interaction networks of endothelial cells subsets with pericytes in SSc BVOs exposed to SSc serum

Since ECs-PEs interactions are critical for physiological vessel development and maturation (33, 34), we wondered whether these interactions might be impaired by exposure of SSc BVOs to SSc_aDU serum. To evaluate this, we leveraged the spatial resolution of CODEX to examine the changes in spatial proximity between BEC subpopulations and PEs in SSc BVOs exposed to serum from different donor groups (Fig. 5A). Upon exposure to healthy serum, PEs were in close spatial proximity to multiple BEC subpopulations, including TIE2 BECs, HEY1 CD34 DLL4 BECs and NR2F2 CD34 BECs (Fig. 5A, B). However, in SSc BVOs exposed to SSc_aDU serum, the interaction patterns were altered: the TIE2 BECs and the HEY1 CD34 DLL4 BECs lost interactions with PEs, while the aSMA NR2F2 BECs and the αSMA DLL4 NR2F2 BECs were spatially avoiding PEs (Fig. 5B). Thus, BEC subsets involved in angiogenesis lost interactions with PEs that are critical for vessel formation, which likely contributes to the angiogenic defects, whereas BEC subsets undergoing EndMT avoided pericytes, likely because the mesenchymal phenotype they acquired allowed them to migrate away from vessels. Of note, the interaction network of SSc BVOs exposed to SSc_noDU serum resembled that of SSc BVOs exposed to healthy serum (Fig. 5B), highlighting on an additional experimental level that BVOs are specifically reacting to vasoactive mediators from SSc patients with digital ulcers rather than to factors associated with SSc in general. These results demonstrate that SSc_aDU serum disrupts the physiologic ECs-PEs interaction networks.

**Figure 5.**
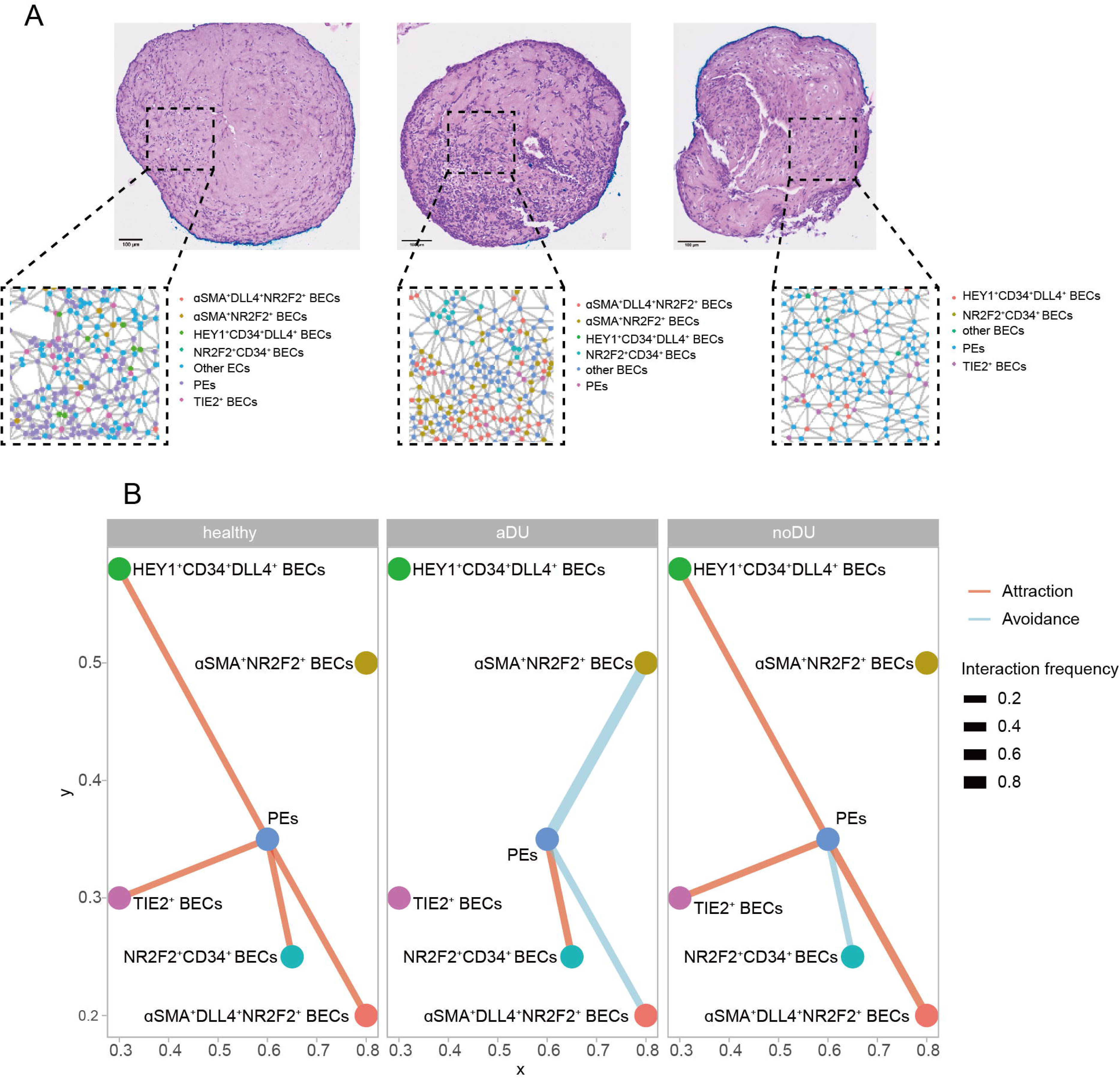
Altered interaction networks between endothelial cell subsets and pericytes in SSc BVOs exposed to SSc serum. (**A**) Representative histological images (HE stainings) and CODEX-based identification of BEC subpopulations and PEs neighboring each other (found in close spatial proximity), exposed to healthy serum, SSc_aDU serum and SSc_noDU serum in SSc BVOs. Scale bars: 100 μm. (**B**) Interaction network analysis of PEs and BEC subpopulations exposed to healthy serum, SSc_aDU serum and SSc_noDU serum in SSc BVOs. The width of the lines represents the interaction frequencies, i.e. proportion of samples exhibiting statistically significant interactions, with red indicating attraction and blue indicating avoidance. SSc: systemic sclerosis; BVO: blood vessel organoids; HE: hematoxilin and eosin; CODEX: co-detection by indexing; BEC: blood endothelial cells; PE: pericytes; SSc_aDU serum: serum from SSc patients with active digital ulcers; SSc_noDU serum: serum from SSc patients without active digital ulcers.

### SSc serum-induced vasculopathy in SSc BVOs is IgG dependent

Since IgG autoantibodies with direct pathogenic roles in SSc vasculopathy were previously described (35–37), we reasoned that IgGs might mediate the deleterious effects of SSc_aDU serum in SSc BVOs. To evaluate this, we treated SSc BVOs with SSc_aDU serum depleted of IgGs or with healthy serum enriched in IgGs from SSc_aDU serum (Fig. 6A, Supplementary Fig. 6).

**Figure 6.**
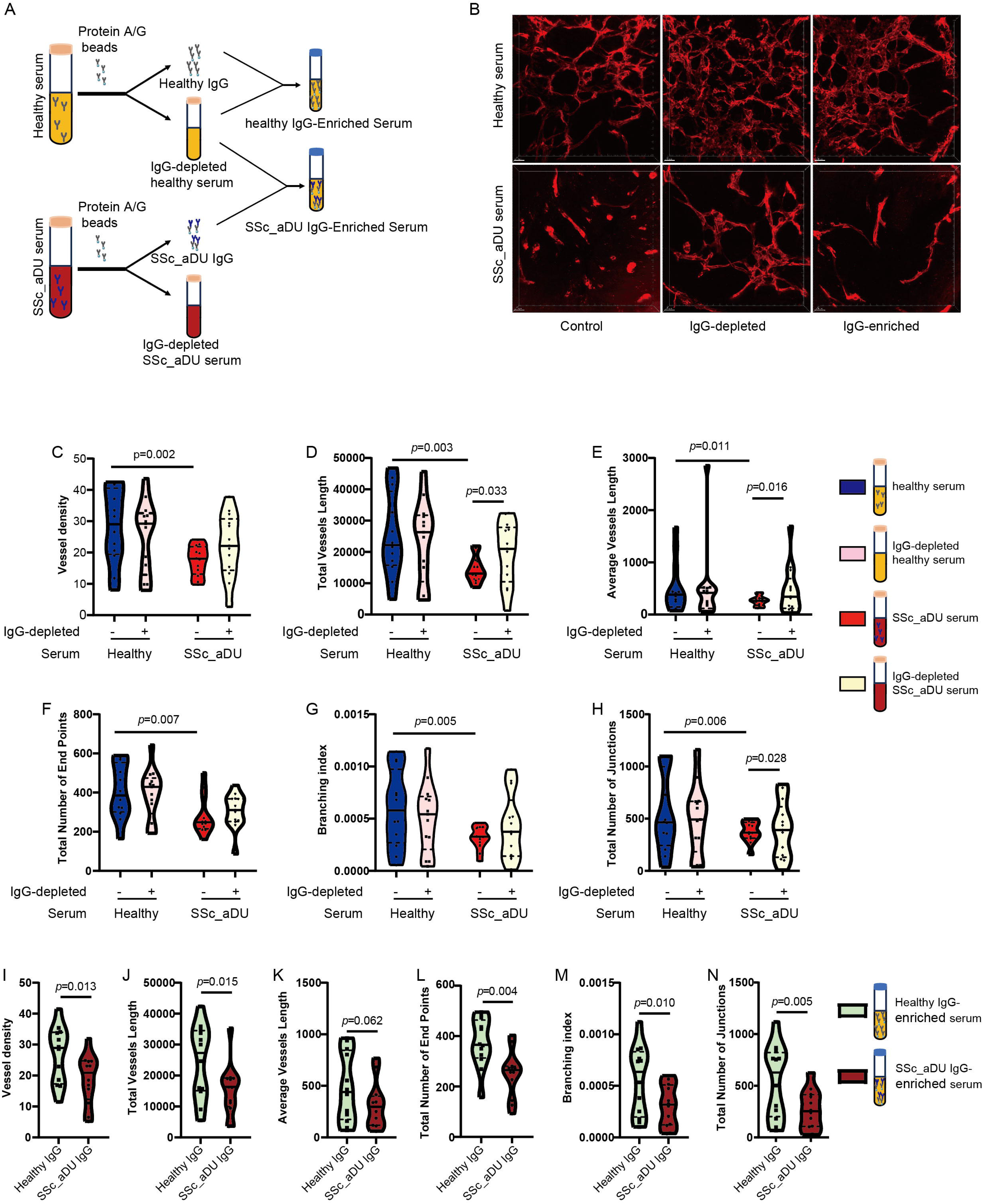
IgG dependency of SSc serum-induced vasculopathy in SSc BVOs. (**A**) Schematic illustration of depletion of IgG from healthy and SSc_aDU serum, and enrichment of IgG-depleted healthy serum with healthy-or SSc_aDU IgG. (**B**) Representative 3D rendering of confocal images of vascular networks in SSc BVOs treated with complete or IgG-depleted healthy serum or SSc_aDU serum, as well as with SSc_aDU- or healthy IgG-enriched healthy serum (after prior IgG depletion). Endothelial cells forming vessel structures were identified by CD31 marker expression (red). Scale bars, 30 μm. (**C-H**) Quantitative analysis of vascular parameters of SSc BVOs exposed to complete or IgG-depleted healthy serum or SSc_aDU serum. (**I-N**) Quantitative analysis of vascular parameters in endothelial cultures treated with healthy IgG-enriched or SSc_aDU IgG-enriched healthy serum (after prior IgG depletion). The analyzed vascular parameters include vessel density (C, I), total vessel length (D, J), average vessel length (E, K), total number of endpoints (F, L), branching index (G, M), and total number of junctions (H, N), shown as violin plots. Each condition included 3 different clones with 4-5 organoids each. *P*-values are included from LMMs with serum treatment as a fixed effect and random intercepts fitted for each cell clone identity. SSc: systemic sclerosis; BVO: blood vessel organoids; IgG: immunoglobulin G; 3D: three-dimensional; SSc_aDU serum: serum from SSc patients with active digital ulcers; SSc_noDU serum: serum from patients without active digital ulcers; LMM: linear mixed-effects model.

IgG depletion from SSc_aDU serum partially restored vascular integrity in SSc BVOs, leading to improved network formation and branching, with increased vessel length and number of junctions and numerical increases in vessel density and branching index compared with SSc BVOs exposed to non-IgG depleted SSc_aDU serum (Fig. 6B-H). Enrichment of IgG-depleted healthy serum with IgGs from SSc_aDU serum phenocopied the effects of SSc_aDU serum, and reduced vessel density, vessel length, branching index and number of junctions in SSc BVOs (Fig. 6B, I-N). These findings highlight that IgGs from SSc patients with severe vasculopathy are central mediators of the effects of SSc_aDU serum in SSc BVOs.

### SSc BVOs exposed to SSc serum as a model for testing of drugs that target SSc vasculopathy

We next aimed to evaluate the potential of using SSc BVOs exposed to SSc_aDU serum as a platform for evaluating drugs targeting SSc-related vasculopathy. GSEA analysis of our RNAseq data identified the endothelin pathway as enriched in SSc BVOs upon exposure to SSc_aDU serum (Fig. 7A). Given that the endothelin receptor antagonist Bosentan (BST) is currently used in clinical practice to prevent the formation of ischemic fingertip ulcers in SSc (38), we aimed to evaluate whether treatment with BST can protect the SSc BVOs from the deleterious effects of SSc_aDU serum. SSc BVOs exposed to SSc_aDU serum and treated with BST had a significantly increased vessel density and total vessel length, numerical differences with increased number of junctions and branching index, and reduced αSMA expression compared to vehicle-treated BVOs (Fig. 7B-H, Q and R), thus demonstrating that BST treatment can partially protect against EndMT and vessel damage induced by SSc serum. We further observed that NOTCH signaling pathway-associated genes were upregulated in ECs treated with SSc_aDU serum compared to those treated with healthy serum in SSc BVOs (Fig. 7I). Aberrant NOTCH signaling has previously been implicated in diabetes-induced microangiopathy (23). γ-secretase inhibitors are the most frequently used pharmacological approach to target NOTCH signaling. Treatment with the γ-secretase inhibitor DAPT protected SSc BVOs from the effects of SSc_aDU serum, with increased vessel density, vessel length, number of junctions, branching index and reduced αSMA expression (Fig. 7J-R).

**Figure 7.**
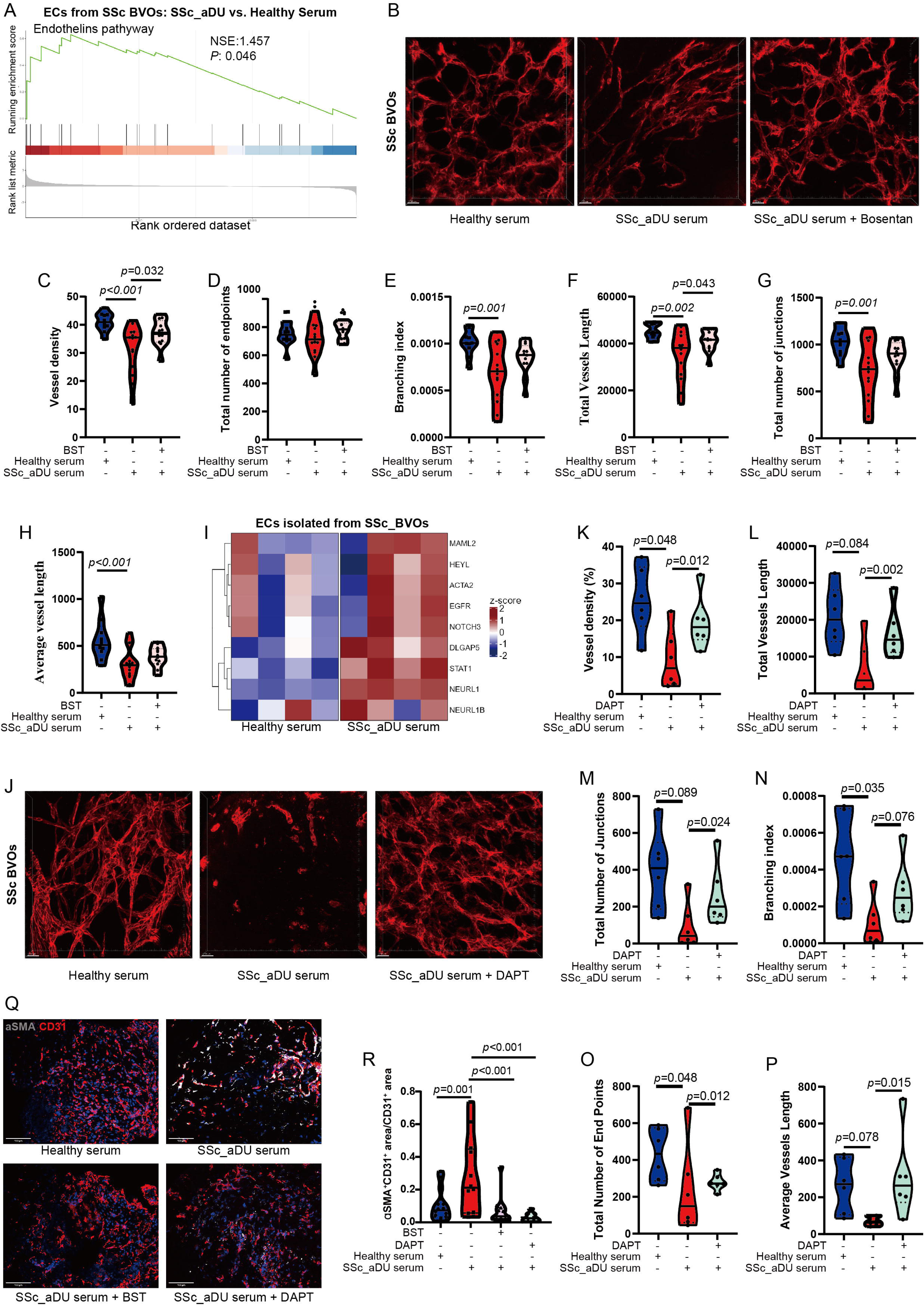
Bosentan and DAPT prevent SSc serum-induced angiogenic defects in SSc BVOs. (**A**) GSEA enrichment plot of RNAseq data for the term “Endothelins Pathway” of ECs purified from SSc BVOs exposed to SSc_aDU serum compared to SSc BVOs exposed to healthy serum. (**B**) Representative 3D rendering of confocal images of vascular networks in SSc BVOs treated with healthy serum, SSc_aDU serum, or BST + SSc_aDU serum for 7 days. CD31 immunostaining (red) was used for identifying vessels. Scale bars, 30 μm. (**C-H**) Quantitative analysis of vascular parameters in BVOs treated with healthy serum, SSc_aDU serum, or BST+SSc_aDU serum, including vessel density (C), total number of endpoints (D), branching index (E), total vessel length (F), total number of junctions (G) and average vessel length (H). (**I**) Heatmap of Notch signaling pathway-related genes in ECs purified from SSc BVOs treated with healthy serum or SSc_aDU serum. (**J**) Representative 3D rendering of confocal images of vascular networks in SSc BVOs treated with healthy serum, SSc_aDU serum or DAPT + SSc_aDU serum for 7 days. CD31 immunostaining (red) was used for identifying vessels. Scale bars = 30 µm. (**K-P**) Quantitative analysis of vascular parameters in BVOs treated with healthy serum, SSc_aDU serum, or DAPT+ SSc_aDU, including vessel density (K), total vessel length (L), total number of junctions (M), branching index (N), total number of endpoints (O), and average vessel length (P). (Q) Representative pictures of αSMA positive endothelial cells in SSc BVOs treated with healthy serum, SSc_aDU serum, BST + SSc_aDU serum, or DAPT + SSc_aDU serum for 7 days. Scale bars, 100 μm (S) Quantification of the percentage of αSMA^+^CD31^+^ area from the whole CD31^+^ area in SSc BVOs treated with healthy serum, SSc_aDU serum, DAPT + SSc_aDU serum, or BST + SSc_aDU serum. Each condition included four iPSC clones with 1–4 organoids each. *P*-values are included from LMMs with serum/BST/DAPT treatment as a fixed effect and random intercepts fitted for each cell clone identity. SSc: systemic sclerosis; BVO: blood vessel organoids; GSEA: Gene Set Enrichment Analysis; RNAseq: RNA sequencing; EC: endothelial cells; SSc_aDU serum: serum from SSc patients with active digital ulcers; 3D: three-dimensional; BST: bosentan; LMM: linear mixed-effects model.

Taken together, these data demonstrate that SSc BVOs exposed to SSc serum from patients with severe vasculopathy can be used for testing of drugs that target SSc vasculopathy.

## Discussion

We demonstrate in the current study that human BVOs can be employed as an *in vitro* model for the study of vascular pathology in SSc and for testing of drugs that target microvasculopathy.

BVOs are differentiated from iPSC to form endothelial cells assembled as three-dimensional vessels residing on a basement membrane and surrounded by pericytes, thus reproducing key features of adult human vessels and being ideally positioned for the study of microvascular responses (23). We show that BVOs recapitulate important endothelial cell populations observed in humans, including lymphatic endothelial cells, endothelial tip and stalk cells, involved in angiogenesis, endothelial progenitor cells, endothelial cells committed to an arterial or venous phenotype and endothelial cells undergoing EndMT. Furthermore, BVOs recapitulate pericyte subpopulations observed in humans and their interaction with endothelial cells.

BVOs maintain the genetic background of the donor of origin and allow to disentangle the relative contribution of genetic and environmental factors to a pathologic vascular phenotype. We demonstrate that BVOs generated from SSc donors, thus maintaining the genetic susceptibility of SSc patients, intrinsically recapitulate early stages of vasculopathy observed in SSc patients, with uncontrolled angiogenesis and formation of immature vessels. Moreover, these SSc-derived BVOs also demonstrate a distinct reaction to external factors compared to healthy BVOs. When exposed to serum from SSc patients with clinically manifest severe microvasculopathy, they recapitulate typical features of vasculopathy in later stages of SSc such as loss of small vessels and presence of avascular areas. These phenotypes observed in SSc-derived BVOs resemble the pathologic vascular changes observed in SSc patients upon evaluation by capillaroscopy as a standard diagnostic tool (8, 9). Notably, SSc serum did not impair angiogenesis in healthy blood vessel organoids, most likely due to the lack of genetic susceptibility. We thus provide, to our knowledge, the first experimental proof for the paradigm inferred from observational studies that circulating factors induce pathologic SSc features in predisposed individuals.

We further elucidate the mechanisms leading to angiogenic defects in SSc blood vessel organoids exposed to SSc serum. We demonstrate in a multi-omics approach that SSc endothelial cells undergo EndMT in the presence of SSc serum, with increased chromatin accessibility assigned to genes related to a mesenchymal phenotype, and decreased accessibility for genes related to an endothelial phenotype, which leads to a corresponding shift in gene expression and pathway activity and to an increased expression of EndMT promoting transcription factors, markers of myofibroblasts and collagen type 1 production. At a single cell level, this is accompanied by a shift from endothelial stalk or tip cell subsets, as pro-angiogenic subpopulations, to subpopulations that express high levels of αSMA, collagen type 1 alongside TFs involved in EndMT. Of note, not only endothelial cells, but also pericytes transition from a PDGFRβ^hi^αSMA^-^ or CD90^hi^ αSMA^-^ phenotype to an αSMA^+^SM22^+^ phenotype and thus contribute to the pool of myofibroblasts. EndMT and pericyte-to-myofibroblast transition have been previously described in the skin of SSc patients and lungs of patients with SSc-ILD (27, 31), thus demonstrating that SSc-BVOs can recapitulate key features of SSc pathology in affected tissues. Not only the fate transitions of endothelial cells and pericytes contribute to the angiogenic defects in SSc blood vessel organoids, but also an altered endothelial cell-pericyte interaction network, with e.g. loss of interactions between TIE2^+^ endothelial stalk cells with pericytes, interactions that are critical for maturation of newly formed vessels (39, 40).

Our data demonstrates that exposure of SSc blood vessel organoids to SSc serum induces a dysregulation of a transcription factor network, with reduced expression and/or activity of transcription factors involved in angiogenesis, such as ETS1, FOXO4 or ELK4, and increased expression and/or activity of transcription factors involved in EndMT, such as ZEB1, SNAI1, SNAI2 or TWIST1. This dysregulation might be the consequence of a sequence of events that involve chromatin remodeling with subsequent changes in mRNA and protein expression to induce transition from an endothelial to a mesenchymal fate.

Thus, exposure of SSc blood vessel organoids to SSc serum induces key pathogenic vascular changes observed in SSc patients that link vasculopathy with tissue fibrosis. Blood vessel organoids might thus be used to study not only vasculopathy, but also the transition and the relative contribution of vasculopathy to tissue fibrosis. Furthermore, our data suggests that endothelial cells from SSc blood vessel organoids exposed to SSc serum upregulate the adhesion molecules ICAM and might promote leukocyte migration and activation. A previous study demonstrated that IL-1β-activated endothelial cells can trigger alternative activation of macrophages, which subsequently induce fibroblast activation (41). Macrophages could be included in blood vessel organoids to study and potentially interfere with the crosstalk endothelial cells-macrophages.

The inhibitory effects of serum from SSc patients with severe SSc vasculopathy are at least in part mediated by IgG autoantibodies, since depletion of IgGs prevents these effects, whereas enrichment of IgG-depleted healthy serum with SSc IgGs phenocopies the effects of complete SSc serum. Several autoantibodies present in SSc patients could potentially mediate these effects. SSc patients with digital ulcers exhibit higher levels of anti-endothelial cell antibodies (AECA) in serum compared to those without digital ulcers (37, 42), and AECA has been identified as a marker of vascular damage (43). Furthermore, anti-AT_1_R and anti-ET_A_R antibody positive IgGs from SSc sera were previously shown to induce changes in the phenotype of endothelial cells (36). However, pathogenic autoantibodies might not be the only mechanism responsible for the deleterious effects of SSc serum. MMP-12-dependent uPAR-D1 cleavage, e.g., was previously shown to mediate the induction of EndMT by SSc serum (44). We further demonstrate that treatment with the ET-1 antagonist BST or with the γ-secretase inhibitor DAPT prevents microvascular changes and EndMT induced by SSc serum in SSc blood vessel organoids. BST was previously shown to prevent angiogenic defects and EndMT in endothelial cells co-cultured with SSc fibroblasts (45), and is in current clinical use in SSc for prevention of recurring digital ulcers (46). The efficacy of BST in BVOs thus serves as a proof of concept that blood vessel organoids respond to well-established drugs that target SSc vasculopathy. Notch signaling was previously shown to be upregulated in SSc (29, 47, 48), and one of the αSMA^+^ endothelial cell subsets defined by our CODEX approach expresses high levels of the Notch ligand DLL4. The efficacy of DAPT in ameliorating SSc serum-induced vasculopathy in SSc BVOs provides evidence that aberrant Notch signaling plays critical roles in mediating angiogenic defects and EndMT in SSc. Our data could thus stimulate follow-up studies with γ-secretase inhibitors for the treatment of SSc-vasculopathy.

In summary, we characterized SSc-BVOs as a novel human model system to study SSc-associated vasculopathy. We demonstrate that genetic predisposition as well as circulating mediators such as autoantibodies from SSc serum synergize to unleash the full spectrum of vascular changes in SSc such as upregulated EndMT, pericyte-to-mesenchymal transition and loss of ECs-PEs interactions in BVOs, which result in dysfunctional angiogenesis and perivascular fibrosis. Exposure of SSc-BVOs to SSc serum triggers changes on epigenetic, mRNA and protein levels to downregulate pro-angiogenic pathways and activate EndMT programs. We also provide evidence that SSc-BVOs can serve as a preclinical model system to evaluate the effects of drug candidates on SSc microvasculopathy and use this model to provide first evidence for potential therapeutic effects of γ-secretase inhibition on SSc vasculopathy.

## Resource availability

RNA-seq data are publicly available at GEO from the date of publication. This paper does not report original code.

Any additional information required to re-analyze the data reported in this paper is available from the lead contact upon request.

## Acknowledgements

We thank Sonja Plötz, Michaela Farrell, Christoph Liebel, Philipp Steinbrecher, Lukas Sokolowski and Wolfgang Espach for excellent technical assistance. We further thank Figdraw for their assistance in creating symbols.

Confocal spinning disc microscopy was enabled on a Zeiss Spinning Disc Axio Observer Z1, funded by Deutsche Forschungsgemeinschaft (DFG, German Research Foundation) — project 248122450. Fluorescence microscopy was performed at the Advanced Light Microscopy Core Facility (Ad-Light), Medical Faculty of the Heinrich-Heine-University, Düsseldorf.

The project was supported by the following grants: Grants DI 1537/17-1, DI 1537/20-1, DI 1537/22-1, DI 1537/23-1, DI 1537/27-1, DI 1537/28-1 of the German Research Foundation (JHWD), MA 9219/2-1 of the German Research Foundation (AEM), SFB CRC1181 (project C01) of the German Research Foundation, grants 2021_EKEA.03 (AEM) and 2022_EKMS.02 (AEM) of the Else-Kröner-Fresenius-Foundation, The Edith Busch and World Scleroderma Foundation Research Grant Programme 2022-2023 (AEM), the German Federal Ministry of Education and Research (BMBF), MASCARA program, TP 2 (01EC1903A), ACS_iIMMUNE (01EO2105 to MR), the Research Committee of the Medical Faculty of the Heinrich-Heine University Düsseldorf (Forschungskommission; ID 2022-18, ID 2023-33 and ID 2023-31 to AHG, AEM and JHWD, respectively), an unrestricted research grant from the Hiller-Foundation (JHWD) and a Career Support Award of Medicine of the Ernst Jung Foundation (JHWD).

Our analysis was supported by a de.NBI Cloud project (YNL) within the German Network for Bioinformatics Infrastructure (de.NBI) and ELIXIR-DE (Forschungszentrum Jülich and W-de.NBI-001, W-de.NBI-004, W-de.NBI-008, W-de.NBI-010, W-de.NBI-013, W-de.NBI-014, W-de.NBI-016, W-de.NBI-022).

## Author contributions

YX, JHWD and AEM designed the study. YX, XH, LX, YNL, BG, SK, AHG, TF, PMB, PT and AEM were involved in acquisition and analysis of data. All authors were involved in interpretation of data. MR, FM, JA and JW provided essential samples. All authors were involved in manuscript preparation and proof-reading.

## Supplementary information

**Document S1**. Figures S1–S6 and Table S1-S2

**Video S1**: Rotational view of 3D reconstructed vessels in BVOs, 360-degree rotational view of a representative 3D reconstruction of CD31^+^ vessel structures in BVOs. BVO: blood vessel organoids; 3D: three-dimensional. Related to Figure 1B

**Video S2**: Rotational view of 3D reconstructed vessels and perivascular structures in BVOs. 360-degree rotational view of a representative 3D reconstruction of endothelial cells (CD31, red), pericytes (PDGFRβ, green), and basement membrane (COLIV, purple) in BVOs. BVO: blood vessel organoids; 3D: three-dimensional. Related to Figure 1C.

## STAR Methods

### Key resources table

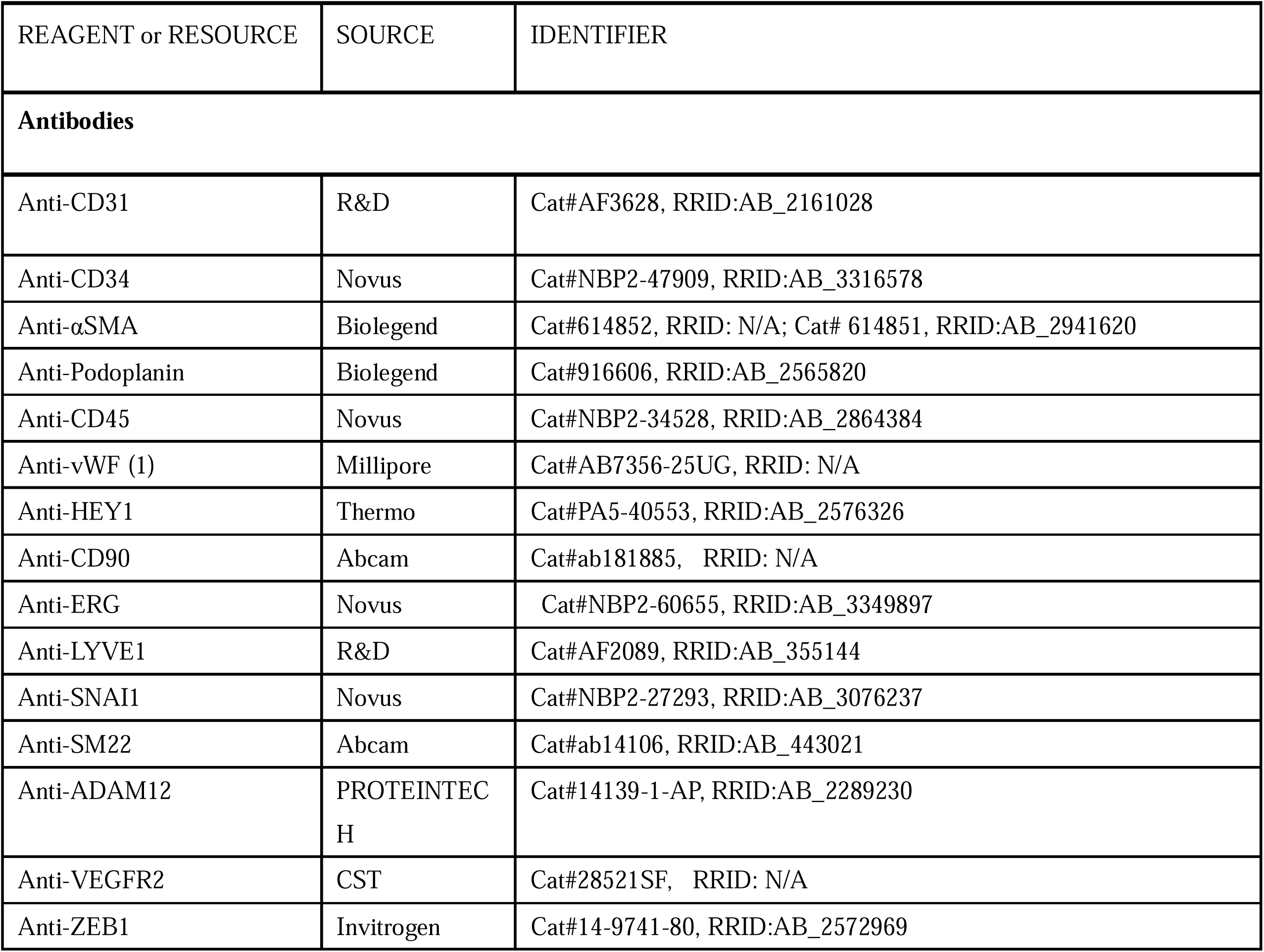
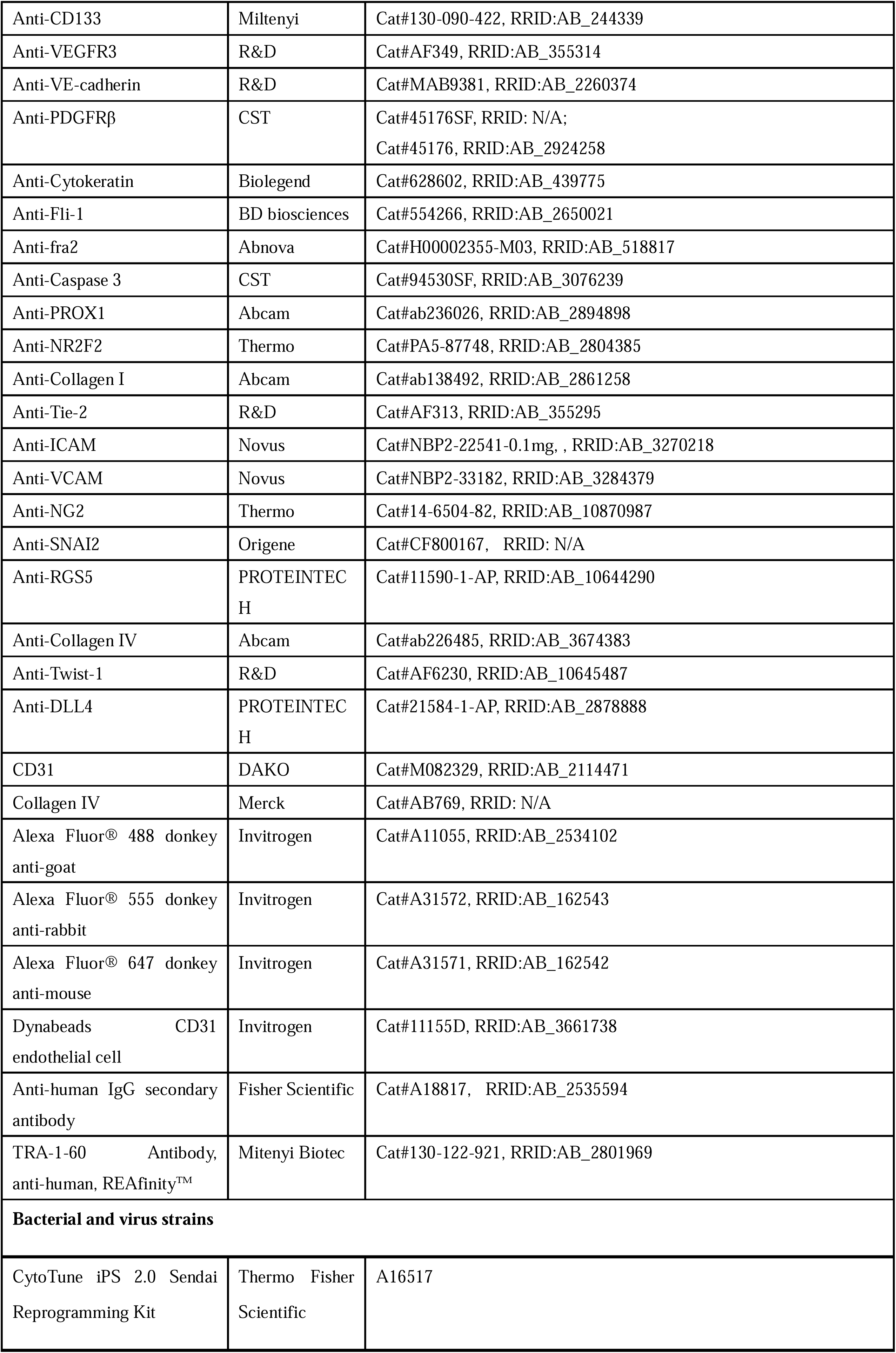
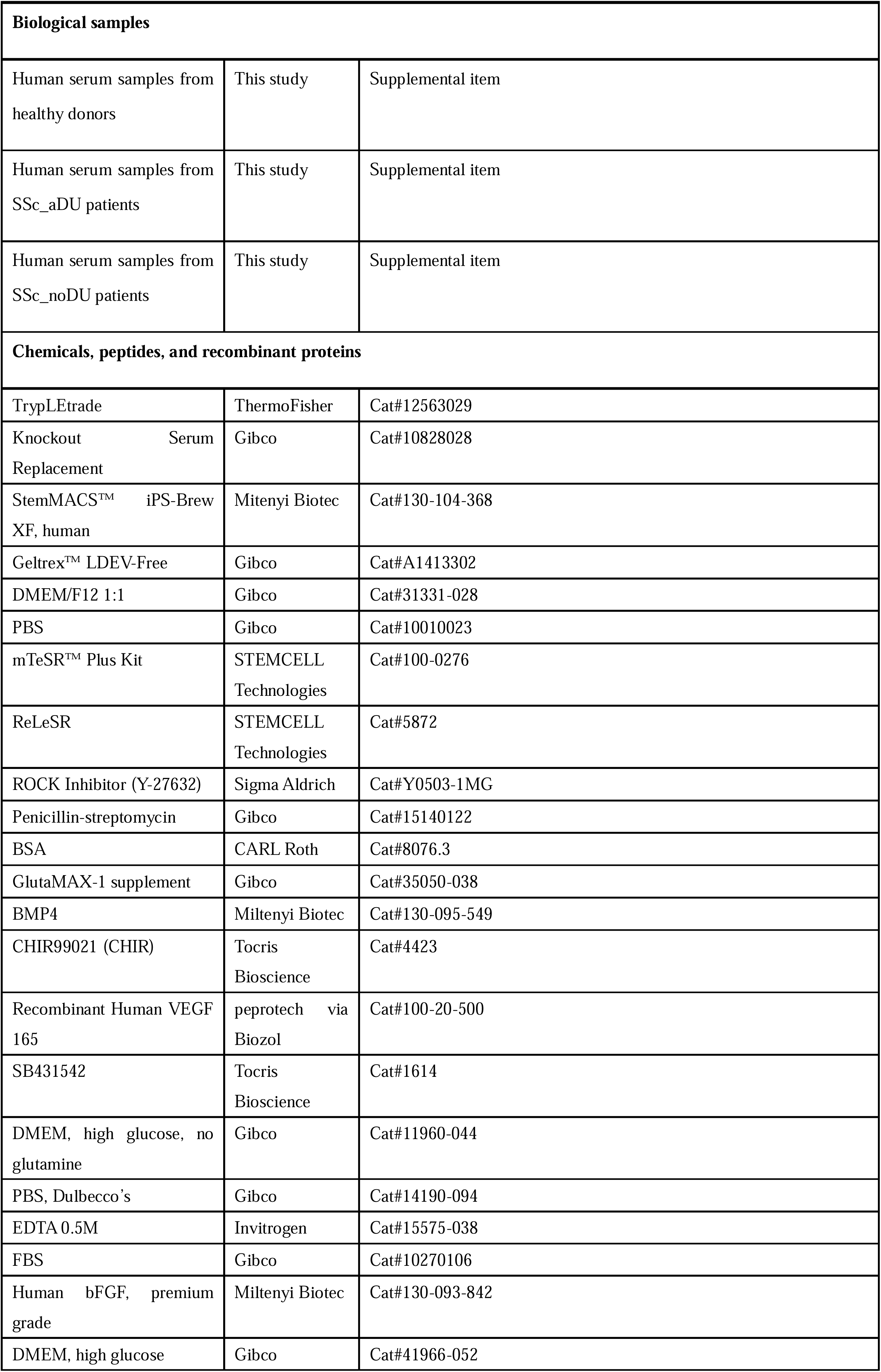
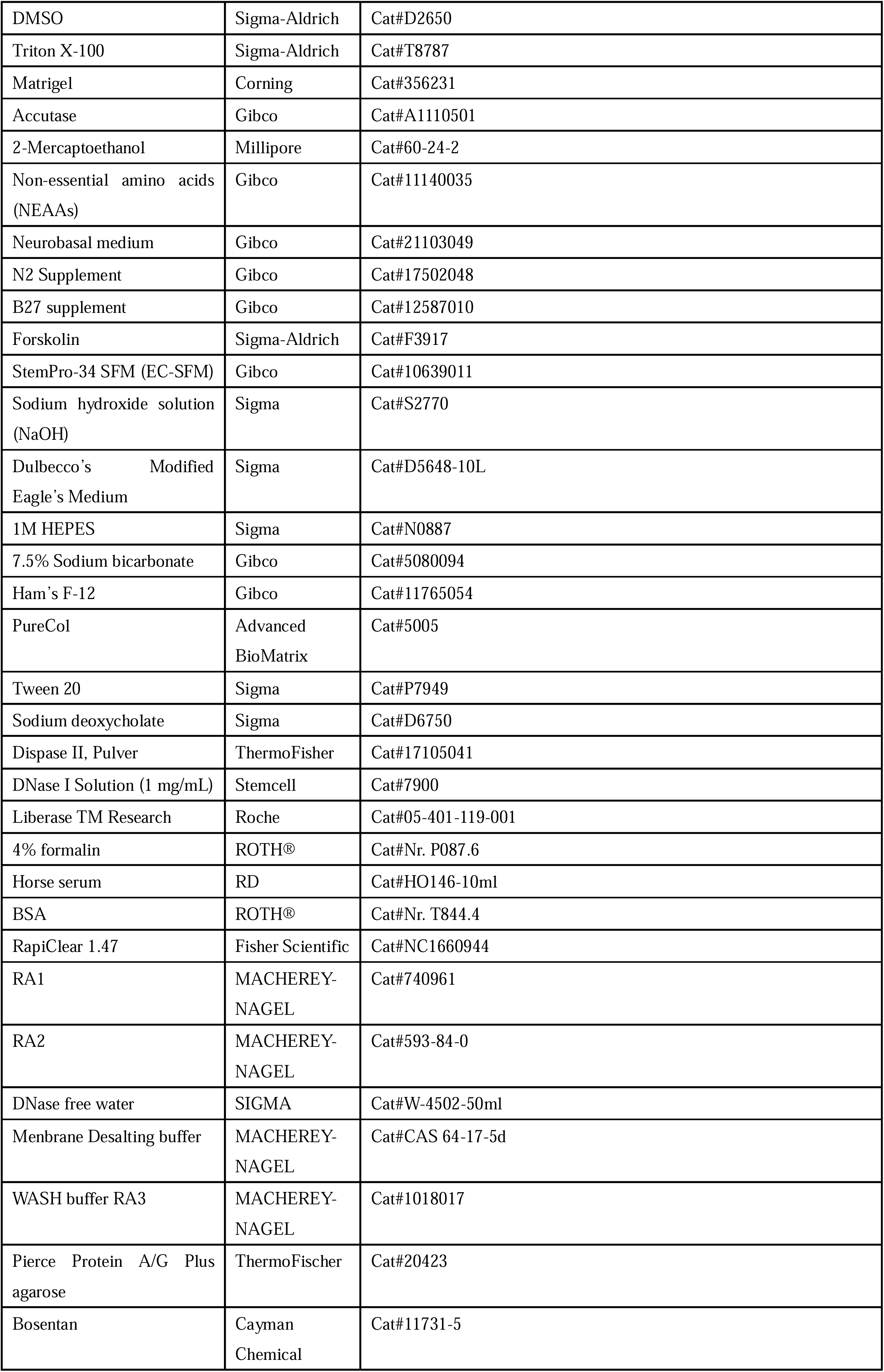
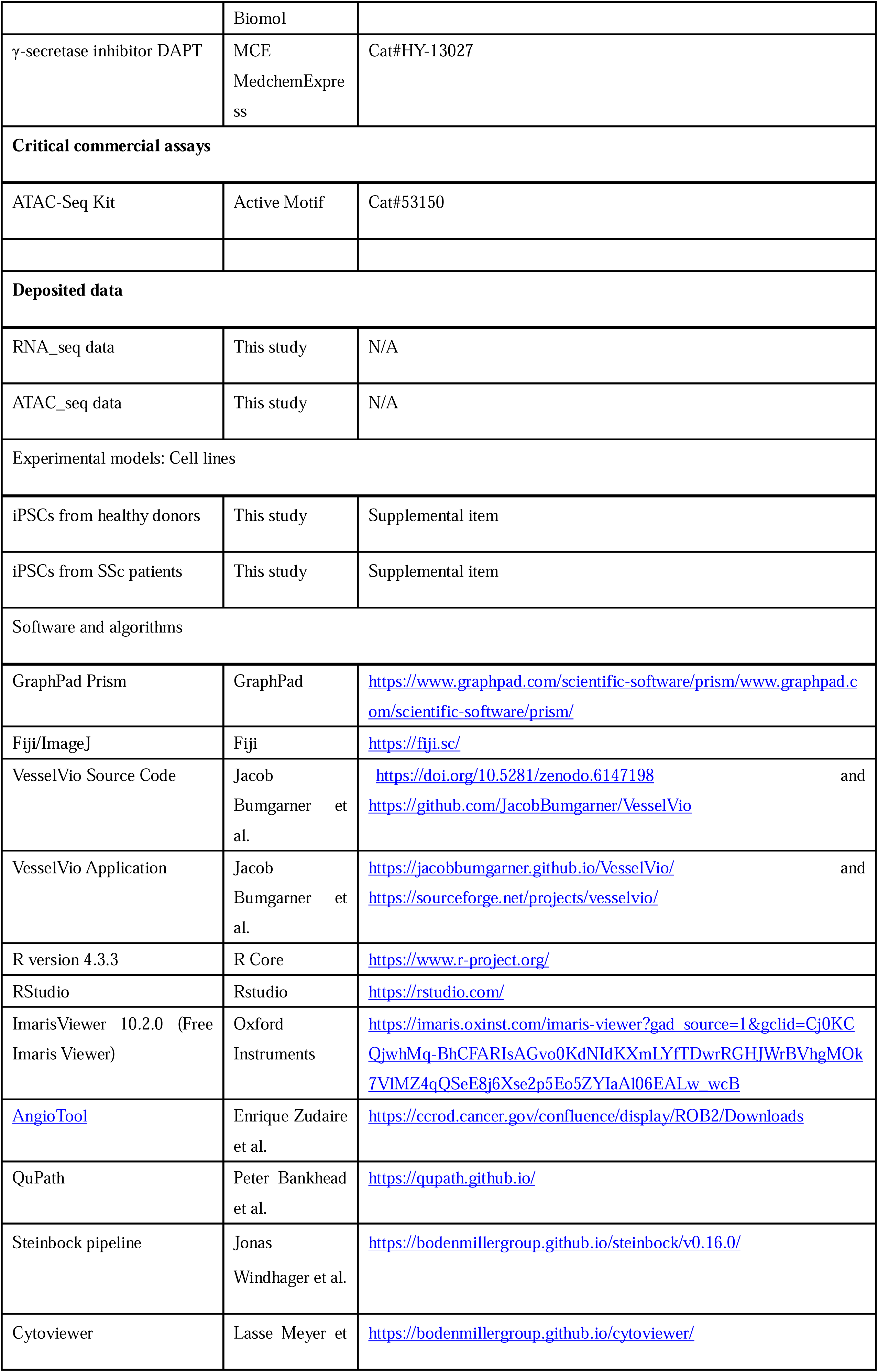
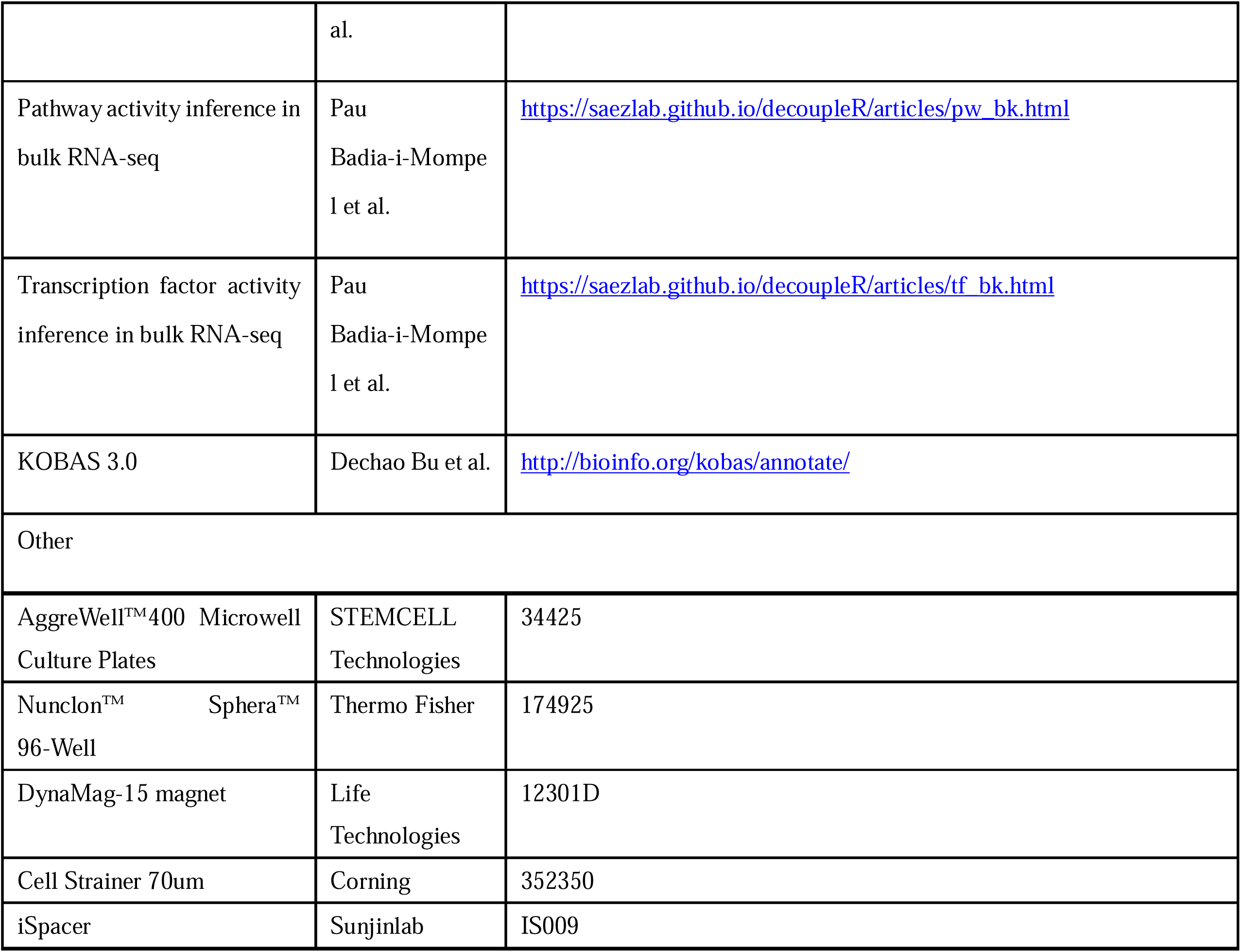

## RESOURCE AVAILABILITY

### Lead contact

Further information, requests for resources, and reagents should be directed to and will be fulfilled by the lead contact, Alexandru-Emil Matei.

### Materials availability

No new reagents were generated specifically for this study. Data and code availability RNA-seq data are publicly available at GEO from the date of publication. This paper does not report original code.

## EXPERIMENTAL MODEL AND STUDY PARTICIPANT DETAILS

### Cell lines

The induced pluripotent stem cell (iPSC) lines used in this study comprised two distinct groups: two healthy control lines and two systemic sclerosis (SSc) patient-derived lines (donor information detailed in Supplementary Table S1). Each iPSC line was clonally expanded to establish two biologically independent subclones, resulting in eight total clones. All cell lines were derived from dermal fibroblasts obtained through punch biopsies. The cell lines were maintained under feeder-free culture conditions for minimum 10 passages prior to experimental use. Regular quality control assessments confirmed absence of mycoplasma contamination in all cell stocks.

### Ethical approval

All donors provided informed consent for participation in the study. Fibroblast isolation and reprogramming to iPSC, as well as the experiments performed with the iPSC were approved by the Ethical Committee of the University of Erlangen-Nürnberg and the Ethical Committee of the University of Düsseldorf (approval numbers: 21-485-Bp, 98_18 B, 30_19B (Erlangen) and 2022-2158 (Düsseldorf)).

## METHOD DETAILS

### Fibroblast reprogramming to iPSCs

Fibroblast reprogramming to iPSCc was performed as previously described (49, 50). Human dermal fibroblasts isolated from skin biopsies of SSc patients or healthy individuals (Table S1) in passages three or four were cultured in six-well plates with DMEM/F12 supplemented with 10% FBS. iPSCs were generated from fibroblasts using the CytoTune iPS 2.0 Sendai Reprogramming Kit (Thermo Fisher Scientific, A16517) according to the manufacturer’s instructions. Briefly, cell lines were transduced with non-integrating Sendai virus containing four reprogramming factors c-MYC, KLF4, OCT3/4, and SOX2. When cell cultures reached 70–80% confluency, iPSCs were washed once with DMEM/F12 (GibcoTM, 31331-028) and incubated with Gentle Cell Dissociation Reagent ReLeSR (STEMCELL Technologie, 5872) for 5 min at room temperature (RT) for passaging. Gentle Cell Dissociation Reagent was aspirated and StemMACS iPS-Brew XF (Mitenyi Biotec, 130-104-368) supplemented with 100 U/mL penicillin/streptomycin was added. Corning® Cell Lifter was used to detach hiPSCs from the cell culture plate. iPSCs were transferred to a new Geltrex-coated plate. Cells with a TRA1-60 positive rate exceeding 75% were selected for further passage and culture. Two clones were used from each donor.

### Generation of BVOs from iPSCs

The iPSCs were initially cultured on 1% Geltrex-coated (Gibco, A1413302) 6-well plates with reduced growth factors in complete mTeSR Plus (STEMCELL Technologies, 100-0276). For directed BVO differentiation, 3 × 10^6^ single iPSCs were cultured in complete mTeSR Plus media supplemented with the ROCK inhibitor Y-27632 (10 μM, Sigma Aldrich, Y0503-1MG) in AggreWell™400 Microwell Culture Plates (STEMCELL Technologies, 34425) for 1 day to form aggregates (denoted as day 0 of vascular network differentiation). Aggregates were treated with mesoderm induction medium composed of 3 mL of N2B27 medium (composed of 50% DMEM:F12 medium (Gibco, 31331-028), 50% Neurobasal medium (Gibco, 21103049), 2% B27 supplement (Gibco, 12587010), 1% N2 supplement (Gibco, 17502048)L, 1% Glutamax (Gibco, 35050-038), 1% Penicillin-streptomycin (Gibco, 15140122), and 7‰ β-mercaptoethanol (Millipore, 60-24-2)) containing 12 µM CHIR99021 (Tocris Bioscience, 4423) and 30 ng/mL BMP-4 (Miltenyi Biotec, 130-095-549) for three days. Thereafter, the medium was changed to N2B27 medium containing 100 ng/mL VEGF-A (Peprotech, 100-20-500) and 2 µM forskolin (Sigma-Aldrich, F3917). After 5 days, the aggregates were seeded into Matrigel (Corning, 356231)): Collagen I (Advanced BioMatrix, 5005) (1:1) gels and treated with complete StemPro-34 SFM medium (Gibco, 10639011) supplemented with 15% FBS (Gibco, 10270106), 100 ng/mL VEGF-A, and 100 ng/mL FGF-2 (Miltenyi Biotec, 130-093-842) to induce blood vessel sprouting. The medium was changed every 3 days thereafter. Blood vessel networks were formed on day 12. Individual vessel networks were then isolated under a dissection microscope and transferred to low adhesion 96-well plates (Thermo Fisher, 174925) (denoted as day 0 of BVO formation).

### Culture of BVOs and exposure to patient serum

Established BVOs (from day 5 after BVO formation) were cultured in BVO culture medium (complete StemPro-34 SFM medium supplemented with 100ng/ml VEGF and 100ng/ml FGF2) along with 15% pooled serum from 5-7 donors, from the following groups: 1) healthy donors (denoted as healthy serum), 2) SSc patients with active digital ulcers and active disease (defined according to the EUSTAR activity index (51)) (denoted as SSc_aDU serum), and 3) SSc patients with active disease, but without active digital ulcers (denoted as SSc_noDU serum). Incubation with serum was performed for one week, and change of medium with patient serum was exchanged after two and five days, and the BVOs were collected seven days after exposure to serum.

### Depletion of IgG from serum

In a subset of experiments, serum was depleted of total IgGs before stimulation of BVOs. Total IgG depletion from serum was performed using Pierce Protein A/G Plus agarose (ThermoFischer, 20423). We mixed 500 µL of either healthy serum or SSc_aDU serum with an equal volume of 50% agarose gel beads and agitated the mixture at 300 rpm on a shaker. The mixture was pipetted up and down every 15 minutes for three hours at room temperature. After centrifugation at 12000g for 10 minutes, the IgG-depleted supernatant was transferred to a new tube, while the agarose gel, containing the IgG-enriched fraction, was retained. The successful depletion of IgG from serum was confirmed by Western Blot. The IgG-depleted healthy serum and SSc_aDU serum were then used at a concentration of 15% in BVO culture medium. BVOs treated with complete healthy and SSc_aDU serum, without IgG depletion, were included in these experiments as controls. Additionally, BVOs were treated with IgG-enriched fractions derived separately from healthy serum and SSc_aDU serum, each diluted in IgG-depleted healthy serum, to compare the effects of healthy IgG and SSc_aDU IgG.

### Western blot analysis

IgG-depleted supernatant, IgG-enriched agarose gel, and pure serum (20µL each) were separately mixed with 5 µL of loading buffer (Laemmli buffer). Each mixture was heated at 100°C for 10 minutes. Following this, the proteins were separated by SDS-PAGE and transferred onto a polyvinylidene fluoride (PVDF) membrane. The membrane was then incubated with an anti-human IgG secondary antibody (1:5000, Fisher Scientific, A18817). Detection was performed using enhanced chemiluminescence (ECL, GE Healthcare). Western blot images were visualized and captured using the ChemiDoc MP imaging system (Bio-Rad).

### Drug treatment of BVOs

In another subset of experiments, drug treatment of BVOs was performed. For this, BVOs were exposed to BVO culture medium containing 15% SSc_aDU serum in the presence or absence of the following drugs: Bosentan (10 µM, Cayman Chemical Biomol, 11731-5) and the γ-secretase inhibitor DAPT (25 µM, MCE MedchemExpress, HY-13027). The treatment lasted for 7 days, with fresh medium changed every other day.

### Immunofluorescence staining of BVOs

Whole-mount BVOs collected for immunofluorescence staining were fixed in 4% PFA for 1 hour at room temperature (RT). Following fixation, they were permeabilized and blocked using the blocking buffer (47.5mL DPBS (Gibco, 14190-094) with 1% BSA (CARL Roth, 8076.3), 3% FBS (Gibco, 10270106), 0.5% Tween 20 (Sigma-Aldrich, P7949), 0.5% Triton X-100 (Sigma-Aldrich, T8787), and 0.01% sodium deoxycholate solution (Sigma-Aldrich, D6750)) overnight at 4°C. Antibodies against CD31 antibodies (1:100, DAKO, M082329), PDGFRβ (1:100, CST, 45176SF) and Collagen IV (1:200, Merck, AB769) were diluted in blocking buffer and incubated for 1 week at 4°C. Next, BVOs were washed with DPBS-T (0.1% Tween 20) three times, each for at least 30 minutes and then incubated with Alexa Fluor 647 donkey-anti-mouse antibodies (1:200, Invitrogen, A31571), Alexa Fluor® 555 donkey anti-rabbit (1:200, Invitrogen, A31572), Alexa Fluor® 647 donkey anti-goat (1:200, Invitrogen, 11055), and DAPI (1:1000) for another 1 week at 4°C. After incubation, BVOs were cleared using RapiClear 1.47 (Fisher Scientific, NC1660944) for 5 minutes before mounting onto iSpacers 1.0 mm (Fisher Scientific, NC1986379) on coverslips. Whole-mount BVOs were imaged using a Zeiss 880 Laser Scanning Microscope with a 10x or 20x objective. For 3D reconstruction, Z-stack images were acquired as follows: with the 10x objective, optical slices were captured across a total depth of approximately 206 µm, spaced 7 µm apart; with the 20x objective, optical slices were captured across a total depth of approximately 50 µm, spaced 1 µm apart. Vessel quantification was performed using AngioTool with the following settings: vessel diameter (>=5 px) and vessel intensity (20–255 px).

Tissue sections from formalin-fixed, paraffin embedded organoids were deparaffinized in xylene (3 × 10 min) under a fume hood, followed by rehydration through a graded ethanol series: 100% ethanol (2 × 5 min), 95% ethanol (5 min), and 80% ethanol (5 min). After rehydration, the sections were washed three times with PBS for 5 min each. Antigen retrieval was performed using a microwave heating method. Tissue sections were heated in 1× Citrate buffer (pH 6) for 2 min per cycle, followed by heating in 1× Tris-EDTA buffer (pH 9) for 2 min per cycle, with the entire procedure repeated for three cycles. After antigen retrieval, the sections were allowed to cool to room temperature (RT) for 20 min, followed by washing with double-distilled water (ddH O) for 5 min. To block nonspecific binding, the sections were incubated with 5% BSA in PBS for 1 h at RT. Subsequently, the sections were incubated overnight at 4°C with the primary antibody, anti-CD31 (1:200, R&D, AF3628) and anti-αSMA (1:500, Sigma, A5228) were diluted in 2% BSA/PBS. After primary antibody incubation, sections were washed three times with PBS for 5 min each. For fluorescent labeling, the sections were incubated for 1 h at RT with a fluorescent secondary antibody conjugated with DAPI (DAPI 1:1000), Alexa Fluor 647 donkey-anti-mouse antibodies, and Alexa Fluor 488 donkey anti-goat, diluted 1:200 in 2% BSA/PBS. Following secondary antibody incubation, the sections were washed three times with PBS for 5 min each to remove any unbound antibody. Finally, the sections were mounted with a fluorescent mounting medium and covered with a coverslip.

Quantification of αSMA-positive endothelial cells was performed using Fiji by calculating the ratio of the α-SMA CD31 double-positive area to the total CD31 area.

### RNA sequencing – library preparation and data analysis

For RNA sequencing (RNAseq), we performed sequencing on whole BVOs (without prior cell enrichment or depletion) and separately on endothelial cells (ECs) purified from BVOs. To isolate single cells from BVOs treated with healthy serum, SSc_aDU serum, and SSc_noDU serum, the BVOs were mechanically disrupted and cells were disaggregated using a solution containing 3mg/mL Dispase II (Gibco, 17105041), 0.8mg/mL Liberase TM (Roche, 05-401-119-001), and 0.05ug/ml DNase I (Stemcell Tech, 7900) in DPBS for 20 minutes at 37°C while rotating. The reaction was stopped by adding 5 ml of 0.1% BSA DPBS. To isolate ECs, the single cells were then incubated with CD31 magnetic beads (Invitrogen, 11155D) diluted in a 0.1% BSA DPBS solution for 20 minutes. The cells were then placed in a 15 ml FACS tube positioned on a DynaMag-15 magnet (Invitrogen, 12301D) and washed three times using 0.1% BSA DPBS. The whole BVOs or ECs isolated from BVOs were lysed in 350 μl of RNA lysis buffer, consisting of 350 μl RA1 (MACHEREY-NAGEL, 740961) and 3.5 μl β-mercaptoethanol (Millipore, 60-24-2). Subsequently, total RNA extraction was carried out using the NucleoSpin RNA II extraction system (Macherey-Nagel). Sequencing was performed on an Illumina NovaSeq, with a sequencing depth of 20 million reads, yielding a total data output of 6 Gb per sample. Paired-end sequencing was performed with a read length of 150 bp. RNA-seq data were processed using the Galaxy workflow (https://usegalaxy.org). Raw paired-end reads were quantified using Salmon in quasi-mapping mode against the Homo sapiens transcriptome (GRCh38) to directly generate gene-level count matrices for subsequent analysis (52). The R package edgeR was used to calculate the trimmed mean of M values for read count normalization, and ComBat-seq was employed for batch effect correction (53). EdgeR was used to identify differentially expressed genes (DEGs) (54). DEGs were defined based on the following thresholds: FDR < 0.05 and |log2 fold-change| > 0.585. Gene Set Enrichment Analysis (GSEA) was conducted using the ClusterProfiler package with the most recent gene sets from Bader lab (55). Pathways with an adjusted *p*-value < 0.05 were considered statistically significant. Upregulated or downregulated genes were subjected to Gene Ontology (GO) enrichment analysis using KOBAS 3.0 (http://bioinfo.org/kobas/annotate/), and terms with a *p*-value <0.05 were selected to generate bubble plots. Transcription factor activity was inferred using decoupleR (56).

### ATAC sequencing library preparation and data analysis

For ATAC-seq, single cells were isolated from BVOs following the same enzymatic dissociation protocol described above. After isolation, cells were processed for tagmentation, library preparation, and sequencing following standard ATAC-seq protocols outlined in the ATAC-Seq Kit (Active Motif, 53150) Manual (https://www.activemotif.com/documents/2182.pdf) (57). Briefly, 100000 fresh cells were resuspended in cold ATAC lysis buffer, filtered through a 40 µm cell strainer, and centrifuged. The supernatant was carefully removed, and the cell pellet was resuspended in Tagmentation Master Mix, consisting of 2X Tagmentation Buffer, PBS, Digitonin, Tween-20, and assembled transposomes. The mixture was incubated at 37°C for 30 minutes to fragment chromatin and insert sequencing adapters. Following transposition, DNA was purified using DNA Purification Binding Buffer and sodium acetate, ensuring optimal pH for DNA binding. The purified DNA was amplified using i7 and i5 indexed primers, and libraries were cleaned up with SPRI beads. DNA concentration was measured using Qubit, and fragment size distribution was assessed with a Bioanalyzer or TapeStation. Sequencing was performed on an Illumina HiSeq 2500, with a sequencing depth of 200 million reads, yielding a total data output of 60 Gb per sample. Paired-end sequencing was performed with a read length of 150 bp.

The raw reads were pre-processed using the nf-core pipeline for ATACseq (58, 59). Adapter trimming was performed from raw reads using Trim Galore. Adapter-trimmed reads were aligned to the human reference genome (hg38) using BWA. Reads mapping to the mitochondrial genome were filtered out with the default nf-core pipeline settings, and duplicate reads were removed using Picard. Peak calling was performed with MACS2 and peaks identified across all samples were merged to generate a consensus peak set. Tn5 transposase insertion sites and transcription start site (TSS) enrichment scores were evaluated to assess ATAC-seq data quality. The ATAC signal intensity for each peak was determined by subtracting background noise. FeatureCounts was used to quantify the number of reads overlapping each consensus peak and then the peaks were assigned to genes according to the HOMER annotations using the default.gtf file of the hg38 genome. Differential accessibility regions (DARs) were identified using the edgeR package, with significance thresholds set at |log fold change| > 0.585 and FDR < 0.05. GSEA was performed with the fgsea package in R. Pathways with an adjusted *p*-value < 0.05 were considered statistically significant.

Upregulated or downregulated genes were subjected to Gene Ontology (GO) enrichment analysis using KOBAS 3.0 (http://bioinfo.org/kobas/annotate/), and terms with a *p*-value <0.05 were selected to generate bubble plots.

### Co-detection by indexing (CODEX)

#### Antibody conjugation with DNA oligonucleotides

We followed a previously established protocol for antibody conjugation, validation, and titration (60). Initially, 35-50 μg of purified, carrier-free antibody was loaded onto filter columns for preparation. The antibody was then reduced by incubation with 360 μl of TCEP solution (containing 2.5 mM TCEP and 2.5 mM EDTA in PBS) at RT for 30 minutes. To stop the reduction reaction and remove TCEP, the columns were washed twice with Buffer C solution (consisting of 1 mM Tris (pH 7.0), 1 mM Tris (pH 7.5), 0.15 M NaCl, 1 mM EDTA (pH 8.0), and 0.02% (wt/vol) NaN3 in ddH_2_O). Subsequently, an aliquot of DNA oligonucleotides was dissolved in 400 μl of PBS and added to the column containing the reduced antibody. After incubation at RT for 2 hours, excess oligos were eliminated by washing the antibody-oligo conjugate solution with high-salt PBS (containing 0.02% (wt/vol) NaN_3_ and 0.9 M NaCl). Following centrifugation to remove the high-salt PBS, the elution was carried out using 2 μl of CODEX antibody stabilizer solution per 1 μg of antibody.

#### Sample preparation for CODEX

Five µm-thick formalin-fixed paraffin-embedded (FFPE) tissue sections from SSc and healthy BVOs treated with healthy serum, SSc_aDU or SSc_noDU serum were prepared for CODEX. Deparaffinization and hydration of sections were achieved using xylene, followed by a graded series of ethanol solutions. Antigen retrieval was conducted using a Tris-EDTA buffer (pH 9.0) in a pressure cooker for 20 minutes. Subsequently, the sections were placed in a bleaching solution consisting of 25 ml PBS, 4.5 ml of 30% (wt/vol) HO (Sigma-Aldrich: 216763), and 0.8 ml of 1M NaOH, and sandwiched between two high-power LED panels at full power for 45 minutes, a process that was repeated twice. After bleaching, the sections were washed with a Hydration Buffer (10 mM Tris, pH 7.5; 0.02% (w/v) NaN; 0.1% (w/v) Triton X-100; 10 mM MgCl-6HO; 150 mM NaCl) and blocked using Staining Buffer (Akoya Bioscience) for 30 minutes, followed by blocking buffer (Akoya Bioscience) for another 30 minutes. The primary antibody cocktail (Table S2) in Staining Buffer was then added, and the sections were incubated on an orbital shaker overnight at 4°C. Post-incubation, the sections were washed and fixed according to the instructions provided by Akoya Bioscience, Inc (60).

#### CODEX imaging

Multicycle imaging was conducted using the PhenoCycler-Fusion 1.0 system equipped with a 20X air objective from Akoya Bioscience. The process utilizes barcoded reporter oligonucleotides, labeled with ATTO550 and ATTO647 by biomers.net GmbH, which allowed for the capture of up to two protein stains per cycle. Nuclear imaging was consistently performed using DAPI staining in each cycle. The PhenoCycler-Fusion system produced aligned multiplex images in QPTIFF format. Images were then trimmed using QuPath software version 0.4.3. (61, 62)

#### Generation and normalization of single-cell data

Single-cell data extraction and analysis were carried out in accordance with a previously established workflow, employing the Steinbock toolkit (61). Single-cell data was generated using the Steinbock pipeline (Version 0.16.0, https://bodenmillergroup.github.io/steinbock/v0.16.0/). This dataset included the mean signal intensity per cell, spatial coordinates for cellular positioning, and cellular neighbors, defined as all adjacent cells within a 50 μm radius from the cell boundary. Non-cellular objects with areas smaller than 20 pixels or exceeding 500 pixels were filtered out. The imcRtools R package was employed to load the processed data and construct a Spatial Experiment (spe) R object. Subsequently, an inverse hyperbolic sine (arcsinh) transformation with a scaling factor of 1 was applied. Normalization was performed using a z-score calculation, with the entire dataset serving as the reference. The processed data were saved in RDS format. The data was visualized using Cytoviewer (Version 1.50, https://bodenmillergroup.github.io/cytoviewer/) (63).

#### Clustering and cell annotation

Uniform manifold approximation and projection (UMAP) and t-Distributed Stochastic Neighbor Embedding (tSNE) plots were generated using the imcRtools R package, which was also used for unsupervised PhenoGraph clustering with k = 100. Batch effects in CODEX data were corrected using mutual nearest neighbors (MNN) batch correction (64). The z-rescaled expression of the following markers was used for metaclustering and identification of BECs, LECs and PEs: CD31, HEY1, NR2F2, VEGFR2, CD34, TIE2, DLL4, LYVE1, VEGFR3, SM22, αSMA, PDGFRβ, ADAM12, RGS5, CD90. The following markers were used for BEC subclustering: CD31, CD34, HEY1, NR2F2, VEGFR2, TIE2, αSMA, DLL4; and for PE subclustering, the following markers were used: αSMA, ADAM12, PDGFRβ, SM22, RGS5, CD34 and CD90. Initial cluster definitions were based on protein expression patterns and verified on UMAP plots. Similar clusters were merged to generate distinct, non-overlapping cell subpopulations, which were then annotated based on specific marker expressions.

#### Cellular interaction analysis

Spatial proximity between cells was assessed through pairwise interaction analysis using the imcRtools R package, as described (65). Statistical significance was evaluated via permutation testing, where the number of neighboring cells for each subpopulation was compared against a null distribution generated by randomly shuffling cell type labels across 1000 permutations. Interactions with a p-value < 0.05 were considered statistically significant. Increased cell counts indicated cellular attraction, whereas decreased counts suggested cellular avoidance. Visualization of interactions, including dot plots and network representations, was performed using ggplot2 and ggraph in R.

## Statistics

The data are presented as median ± interquartile range (IQR). A linear mixed-effects model (LMM) was employed to compare the effects of different treatments within the same iPSC clone, correcting for between-clone variability, with serum treatment as a fixed effect and random intercepts fitted for each iPSC clone, unless otherwise specified. The comparisons between two groups where different iPSC clones were present in the two groups were conducted using a Mann-Whitney U test. Statistical analysis was performed using GraphPad Prism 9.4.1 and R (4.4). Comparisons with a P < 0.05 were considered statistically significant.

## References

1. Krieg T, Abraham D, Lafyatis R. Fibrosis in connective tissue disease: the role of the myofibroblast and fibroblast-epithelial cell interactions. Arthritis Res Ther. 2007;9 Suppl 2(Suppl 2):S4.

2. Spagnolo P, Cordier JF, Cottin V. Connective tissue diseases, multimorbidity and the ageing lung. Eur Respir J. 2016;47(5):1535–58.

3. Denton CP, Khanna D. Systemic sclerosis. Lancet. 2017;390(10103):1685–99.

4. Distler JHW, Gyorfi AH, Ramanujam M, Whitfield ML, Konigshoff M, Lafyatis R. Shared and distinct mechanisms of fibrosis. Nat Rev Rheumatol. 2019;15(12):705–30.

5. Kawaguchi Y, Kuwana M. Pathogenesis of vasculopathy in systemic sclerosis and its contribution to fibrosis. Curr Opin Rheumatol. 2023;35(6):309–16.

6. Altorok N, Wang Y, Kahaleh B. Endothelial dysfunction in systemic sclerosis. Curr Opin Rheumatol. 2014;26(6):615–20.

7. Matucci-Cerinic M, Kahaleh B, Wigley FM. Review: evidence that systemic sclerosis is a vascular disease. Arthritis and rheumatism. 2013;65(8):1953–62.

8. Cutolo M, Sulli A, Pizzorni C, Accardo S. Nailfold videocapillaroscopy assessment of microvascular damage in systemic sclerosis. J Rheumatol. 2000;27(1):155–60.

9. Lambova SN, Müller-Ladner U. Nailfold capillaroscopy in systemic sclerosis - state of the art: The evolving knowledge about capillaroscopic abnormalities in systemic sclerosis. J Scleroderma Relat Disord. 2019;4(3):200–11.

10. Manetti M, Guiducci S, Ibba-Manneschi L, Matucci-Cerinic M. Mechanisms in the loss of capillaries in systemic sclerosis: angiogenesis versus vasculogenesis. J Cell Mol Med. 2010;14(6a):1241–54.

11. Shu C, Du W, Mao X, Li Y, Zhu Q, Wang W, et al. Possible single-nucleotide polymorphism loci associated with systemic sclerosis susceptibility: a genetic association study in a Chinese Han population. PLoS One. 2014;9(12):e113197.

12. Kawaguchi Y, Tochimoto A, Ichikawa N, Harigai M, Hara M, Kotake S, et al. Association of IL1A gene polymorphisms with susceptibility to and severity of systemic sclerosis in the Japanese population. Arthritis Rheum. 2003;48(1):186–92.

13. Manetti M, Guiducci S, Romano E, Ceccarelli C, Bellando-Randone S, Conforti ML, et al. Overexpression of VEGF165b, an inhibitory splice variant of vascular endothelial growth factor, leads to insufficient angiogenesis in patients with systemic sclerosis. Circ Res. 2011;109(3):e14–26.

14. Moritz F, Schniering J, Distler JHW, Gay RE, Gay S, Distler O, et al. Tie2 as a novel key factor of microangiopathy in systemic sclerosis. Arthritis Res Ther. 2017;19(1):105.

15. Liakouli V, Cipriani P, Marrelli A, Alvaro S, Ruscitti P, Giacomelli R. Angiogenic cytokines and growth factors in systemic sclerosis. Autoimmun Rev. 2011;10(10):590–4.

16. Avouac J, Wipff J, Goldman O, Ruiz B, Couraud PO, Chiocchia G, et al. Angiogenesis in systemic sclerosis: impaired expression of vascular endothelial growth factor receptor 1 in endothelial progenitor-derived cells under hypoxic conditions. Arthritis Rheum. 2008;58(11):3550–61.

17. Pu W, Wu W, Liu Q, Ma Y, Tu W, Zuo X, et al. Exome-Wide Association Analysis Suggests LRP2BP as a Susceptibility Gene for Endothelial Injury in Systemic Sclerosis in the Han Chinese Population. J Invest Dermatol. 2021;141(5):1254–63 e6.

18. Lopez-Isac E, Acosta-Herrera M, Kerick M, Assassi S, Satpathy AT, Granja J, et al. GWAS for systemic sclerosis identifies multiple risk loci and highlights fibrotic and vasculopathy pathways. Nat Commun. 2019;10(1):4955.

19. Takagi K, Kawamoto M, Higuchi T, Tochimoto A, Harigai M, Kawaguchi Y. Single nucleotide polymorphisms of the HIF1A gene are associated with susceptibility to pulmonary arterial hypertension in systemic sclerosis and contribute to SSc-PAH disease severity. Int J Rheum Dis. 2020;23(5):674–80.

20. Maurer B, Distler JH, Distler O. The Fra-2 transgenic mouse model of systemic sclerosis. Vascul Pharmacol. 2013;58(3):194–201.

21. Manetti M, Rosa I, Milia AF, Guiducci S, Carmeliet P, Ibba-Manneschi L, et al. Inactivation of urokinase-type plasminogen activator receptor (uPAR) gene induces dermal and pulmonary fibrosis and peripheral microvasculopathy in mice: a new model of experimental scleroderma? Ann Rheum Dis. 2014;73(9):1700–9.

22. Wimmer RA, Leopoldi A, Aichinger M, Kerjaschki D, Penninger JM. Generation of blood vessel organoids from human pluripotent stem cells. Nat Protoc. 2019;14(11):3082–100.

23. Wimmer RA, Leopoldi A, Aichinger M, Wick N, Hantusch B, Novatchkova M, et al. Human blood vessel organoids as a model of diabetic vasculopathy. Nature. 2019;565(7740):505-10.

24. Zudaire E, Gambardella L, Kurcz C, Vermeren S. A computational tool for quantitative analysis of vascular networks. PLoS One. 2011;6(11):e27385.

25. Bumgarner JR, Nelson RJ. Open-source analysis and visualization of segmented vasculature datasets with VesselVio. Cell Rep Methods. 2022;2(4):100189.

26. Matucci-Cerinic M, Manetti M, Bruni C, Chora I, Bellando-Randone S, Lepri G, et al. The “myth” of loss of angiogenesis in systemic sclerosis: a pivotal early pathogenetic process or just a late unavoidable event? Arthritis Research & Therapy. 2017;19(1):162.

27. Romano E, Rosa I, Fioretto BS, Manetti M. Recent Insights into Cellular and Molecular Mechanisms of Defective Angiogenesis in Systemic Sclerosis. Biomedicines [Internet]. 2024; 12(6).

28. Rius Rigau A, Li YN, Matei AE, Gyorfi AH, Bruch PM, Koziel S, et al. Characterization of Vascular Niche in Systemic Sclerosis by Spatial Proteomics. Circ Res. 2024;134(7):875–91.

29. Huang M, Tabib T, Khanna D, Assassi S, Domsic R, Lafyatis R. Single-cell transcriptomes and chromatin accessibility of endothelial cells unravel transcription factors associated with dysregulated angiogenesis in systemic sclerosis. Ann Rheum Dis. 2024;83(10):1335–44.

30. Ma F, Tsou PS, Gharaee-Kermani M, Plazyo O, Xing X, Kirma J, et al. Systems-based identification of the Hippo pathway for promoting fibrotic mesenchymal differentiation in systemic sclerosis. Nat Commun. 2024;15(1):210.

31. Di Benedetto P, Ruscitti P, Berardicurti O, Vomero M, Navarini L, Dolo V, et al. Endothelial-to-mesenchymal transition in systemic sclerosis. Clin Exp Immunol. 2021;205(1):12–27.

32. Patnaik E, Lyons M, Tran K, Pattanaik D. Endothelial Dysfunction in Systemic Sclerosis. Int J Mol Sci. 2023;24(18).

33. Teichert M, Milde L, Holm A, Stanicek L, Gengenbacher N, Savant S, et al. Pericyte-expressed Tie2 controls angiogenesis and vessel maturation. Nat Commun. 2017;8:16106.

34. Aguilera KY, Brekken RA. Recruitment and retention: factors that affect pericyte migration. Cell Mol Life Sci. 2014;71(2):299–309.

35. Ahmed SS, Tan FK, Arnett FC, Jin L, Geng YJ. Induction of apoptosis and fibrillin 1 expression in human dermal endothelial cells by scleroderma sera containing anti-endothelial cell antibodies. Arthritis Rheum. 2006;54(7):2250–62.

36. Kill A, Tabeling C, Undeutsch R, Kuhl AA, Gunther J, Radic M, et al. Autoantibodies to angiotensin and endothelin receptors in systemic sclerosis induce cellular and systemic events associated with disease pathogenesis. Arthritis Res Ther. 2014;16(1):R29.

37. Kayser C, Fritzler MJ. Autoantibodies in systemic sclerosis: unanswered questions. Front Immunol. 2015;6:167.

38. Sfikakis PP, Papamichael C, Stamatelopoulos KS, Tousoulis D, Fragiadaki KG, Katsichti P, et al. Improvement of vascular endothelial function using the oral endothelin receptor antagonist bosentan in patients with systemic sclerosis. Arthritis Rheum. 2007;56(6):1985–93.

39. van Splunder H, Villacampa P, Martínez-Romero A, Graupera M. Pericytes in the disease spotlight. Trends in Cell Biology. 2024;34(1):58–71.

40. Savant S, La Porta S, Budnik A, Busch K, Hu J, Tisch N, et al. The Orphan Receptor Tie1 Controls Angiogenesis and Vascular Remodeling by Differentially Regulating Tie2 in Tip and Stalk Cells. Cell Reports. 2015;12(11):1761–73.

41. Laurent P, Lapoirie J, Leleu D, Levionnois E, Grenier C, Jurado-Mestre B, et al. Interleukin-1β–Activated Microvascular Endothelial Cells Promote DC-SIGN–Positive Alternatively Activated Macrophages as a Mechanism of Skin Fibrosis in Systemic Sclerosis. Arthritis & Rheumatology. 2022;74(6):1013–26.

42. Sunderkotter C, Herrgott I, Bruckner C, Moinzadeh P, Pfeiffer C, Gerss J, et al. Comparison of patients with and without digital ulcers in systemic sclerosis: detection of possible risk factors. Br J Dermatol. 2009;160(4):835–43.

43. Abraham DJ, Krieg T, Distler J, Distler O. Overview of pathogenesis of systemic sclerosis. Rheumatology (Oxford). 2009;48 Suppl 3:iii3-7.

44. Manetti M, Romano E, Rosa I, Guiducci S, Bellando-Randone S, De Paulis A, et al. Endothelial-to-mesenchymal transition contributes to endothelial dysfunction and dermal fibrosis in systemic sclerosis. Ann Rheum Dis. 2017;76(5):924–34.

45. Corallo C, Cutolo M, Kahaleh B, Pecetti G, Montella A, Chirico C, et al. Bosentan and macitentan prevent the endothelial-to-mesenchymal transition (EndoMT) in systemic sclerosis: in vitro study. Arthritis Res Ther. 2016;18(1):228.

46. Matucci-Cerinic M, Denton CP, Furst DE, Mayes MD, Hsu VM, Carpentier P, et al. Bosentan treatment of digital ulcers related to systemic sclerosis: results from the RAPIDS-2 randomised, double-blind, placebo-controlled trial. Ann Rheum Dis. 2011;70(1):32–8.

47. Dees C, Zerr P, Tomcik M, Beyer C, Horn A, Akhmetshina A, et al. Inhibition of Notch signaling prevents experimental fibrosis and induces regression of established fibrosis. Arthritis Rheum. 2011;63(5):1396–404.

48. Hutchins T, Sanyal A, Esencan D, Lafyatis R, Jacobe H, Torok KS. Characterization of Endothelial Cell Subclusters in Localized Scleroderma Skin with Single-Cell RNA Sequencing Identifies NOTCH Signaling Pathway. Int J Mol Sci. 2024;25(19).

49. Krach F, Stemick J, Boerstler T, Weiss A, Lingos I, Reischl S, et al. An alternative splicing modulator decreases mutant HTT and improves the molecular fingerprint in Huntington’s disease patient neurons. Nature Communications. 2022;13(1):6797.

50. Popp B, Krumbiegel M, Grosch J, Sommer A, Uebe S, Kohl Z, et al. Need for high-resolution Genetic Analysis in iPSC: Results and Lessons from the ForIPS Consortium. Scientific Reports. 2018;8(1):17201.

51. Valentini G, Iudici M, Walker UA, Jaeger VK, Baron M, Carreira P, et al. The European Scleroderma Trials and Research group (EUSTAR) task force for the development of revised activity criteria for systemic sclerosis: derivation and validation of a preliminarily revised EUSTAR activity index. Ann Rheum Dis. 2017;76(1):270–6.

52. Patro R, Duggal G, Love MI, Irizarry RA, Kingsford C. Salmon provides fast and bias-aware quantification of transcript expression. Nat Methods. 2017;14(4):417–9.

53. Zhang Y, Parmigiani G, Johnson WE. ComBat-seq: batch effect adjustment for RNA-seq count data. NAR Genomics and Bioinformatics. 2020;2(3):lqaa078.

54. Robinson MD, McCarthy DJ, Smyth GK. edgeR: a Bioconductor package for differential expression analysis of digital gene expression data. Bioinformatics. 2010;26(1):139–40.

55. Reimand J, Isserlin R, Voisin V, Kucera M, Tannus-Lopes C, Rostamianfar A, et al. Pathway enrichment analysis and visualization of omics data using g:Profiler, GSEA, Cytoscape and EnrichmentMap. Nature Protocols. 2019;14(2):482–517.

56. Badia IMP, Velez Santiago J, Braunger J, Geiss C, Dimitrov D, Muller-Dott S, et al. decoupleR: ensemble of computational methods to infer biological activities from omics data. Bioinform Adv. 2022;2(1):vbac016.

57. Corces MR, Trevino AE, Hamilton EG, Greenside PG, Sinnott-Armstrong NA, Vesuna S, et al. An improved ATAC-seq protocol reduces background and enables interrogation of frozen tissues. Nat Methods. 2017;14(10):959–62.

58. Patel H, Espinosa-Carrasco J, Langer B, Ewels P, Bot N-C, Garcia MU, et al. nf-core/atacseq: [2.1.2] - 2022-08-07. 2.1.2 ed: Zenodo; 2023.

59. Ewels PA, Peltzer A, Fillinger S, Patel H, Alneberg J, Wilm A, et al. The nf-core framework for community-curated bioinformatics pipelines. Nat Biotechnol. 2020;38(3):276–8.

60. Black S, Phillips D, Hickey JW, Kennedy-Darling J, Venkataraaman VG, Samusik N, et al. CODEX multiplexed tissue imaging with DNA-conjugated antibodies. Nat Protoc. 2021;16(8):3802–35.

61. Windhager J, Zanotelli VRT, Schulz D, Meyer L, Daniel M, Bodenmiller B, et al. An end-to-end workflow for multiplexed image processing and analysis. Nat Protoc. 2023;18(11):3565–613.

62. Greenwald NF, Miller G, Moen E, Kong A, Kagel A, Dougherty T, et al. Whole-cell segmentation of tissue images with human-level performance using large-scale data annotation and deep learning. Nat Biotechnol. 2022;40(4):555–65.

63. Meyer L, Eling N, Bodenmiller B. cytoviewer: an R/Bioconductor package for interactive visualization and exploration of highly multiplexed imaging data. BMC Bioinformatics. 2024;25(1):9.

64. Haghverdi L, Lun ATL, Morgan MD, Marioni JC. Batch effects in single-cell RNA-sequencing data are corrected by matching mutual nearest neighbors. Nat Biotechnol. 2018;36(5):421–7.

65. Rius Rigau A, Liang M, Devakumar V, Neelagar R, Matei AE, Gyorfi AH, et al. Imaging mass cytometry-based characterisation of fibroblast subsets and their cellular niches in systemic sclerosis. Ann Rheum Dis. 2024.

