## Supplemental Data 1 for "Human blood vessel organoids recapitulate key mechanisms of transition from vasculopathy to fibrosis in systemic sclerosis"

**Document S1**

Figure S1. Comparison of vessel structures of SSc and healthy BVOs.

Figure S2. Angiogenic defects in SSc BVOs induced by SSc serum from patients with clinically manifest microangiopathy.

Figure S3. SSc_aDU serum promotes EndMT and suppresses angiogenesis in SSc BVOs.

Figure S4. Main populations in BVOs and shifts in endothelial cell subsets in healthy BVOs upon treatment with SSc serum.

Figure S5. CODEX-based characterization of pericyte subpopulations in SSc BVOs and their shifts upon treatment with SSc serum.

Figure S6. Efficiency of IgG depletion from human serum.

Table S1. Demographics of iPSC donors

Table S2. CODEX Antibody Panel

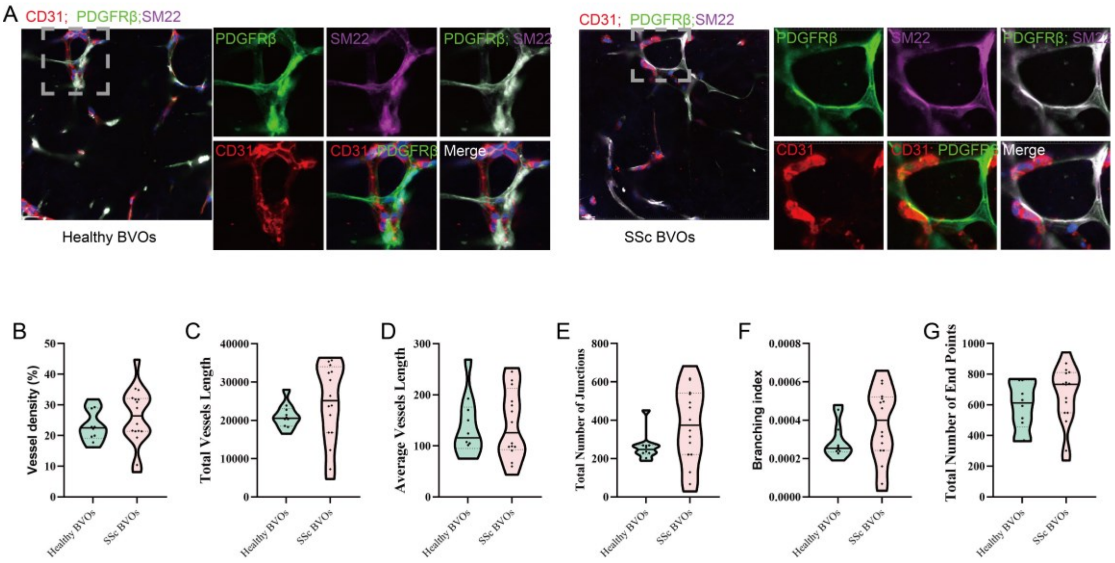


**Figure S1. Comparison of vessel structures of SSc and healthy BVOs.** (A) Representative confocal immunofluorescence images of endothelial cells (CD31$^{+}$, red), pericytes (PDGFRβ$^{+}$, green), and smooth muscle marker (SM22, purple) in both healthy and SSc BVOs (maximum intensity projection). Scale bars: 30 μm. (B-G) Quantitative analysis of vascular parameters between healthy and SSc BVOs using AngioTool, including vessel density (B), total vessel length (C), average vessel length (D), total number of junctions (E), branching index (F), and total number of endpoints (G). Each condition included four different iPSC clones with 2-4 organoids each. Statistically significant p-values (Mann-Whitney U test) are included. SSc: systemic sclerosis; BVO: blood vessel organoids.

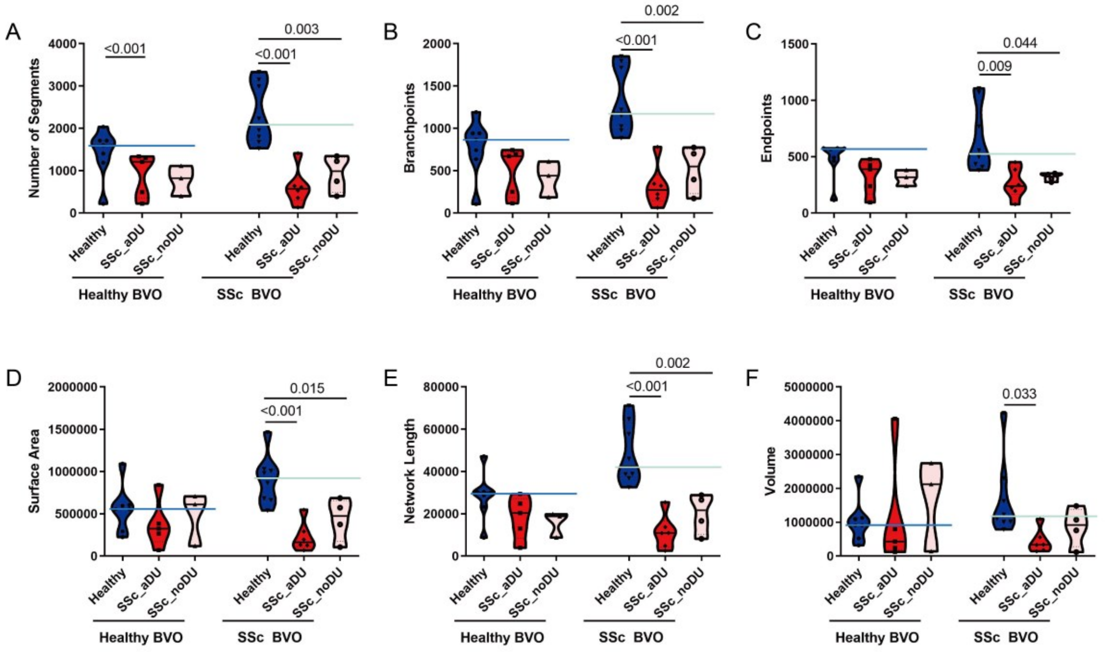


**Figure S2. Angiogenic defects in SSc BVOs induced by SSc serum from patients with clinically manifest microangiopathy.** (A-F) Quantitative analysis of vascular parameters between healthy, SSc_aDU, and SSc_noDU serum using VesselVio and ImageJ, including (A) number of segments, (B) number of branchpoints, (C) number of endpoints, (D) surface area, (E) network length, (F) volume. Each condition includes four iPSC clones with 1–2 organoids each. P-values are included from LMMs with serum treatment as a fixed effect and random intercepts fitted for each cell clone identity. SSc: systemic sclerosis; BVO: blood vessel organoids; 3D: three-dimensional; SSc_aDU serum: serum from SSc patients with active digital ulcers; SSc_noDU serum: serum from patients without active digital ulcers; LMM: linear mixed-effects model.

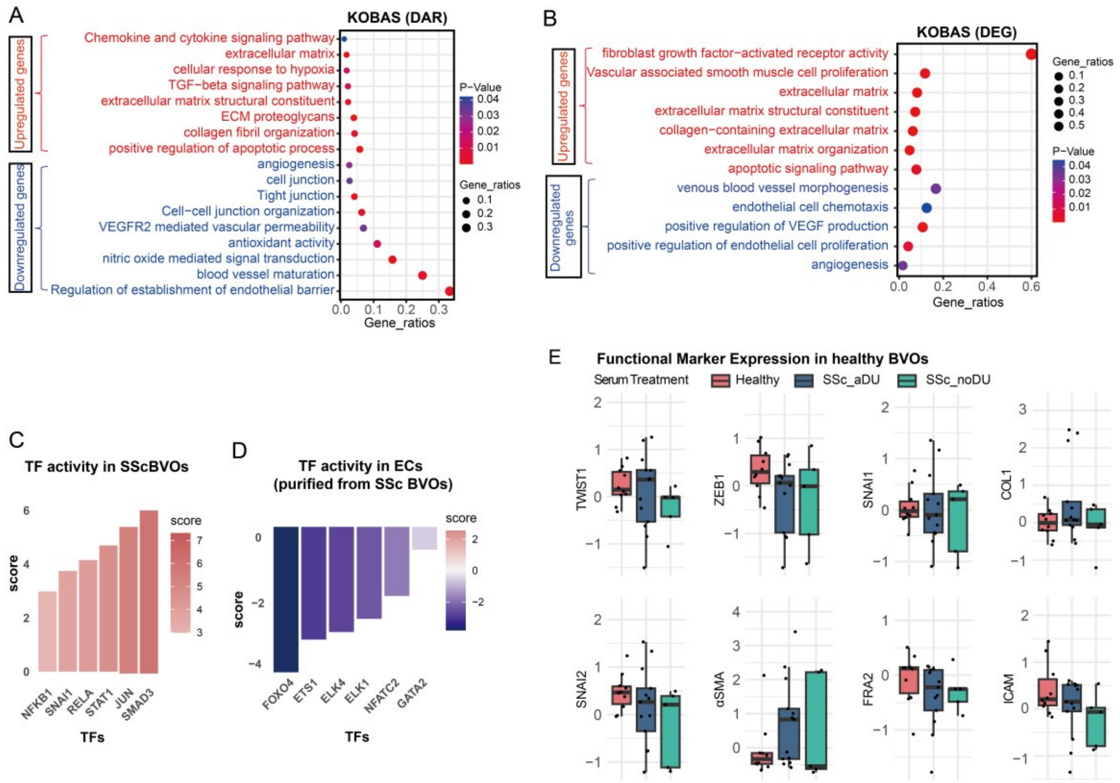

**Figure S3. SSc_aDU serum promotes EndMT and suppresses angiogenesis in SSc BVOs.** (A) Bubble plot of GO pathway enrichment analysis of genes with DARs between SSc BVOs treated with SSc_aDU serum and healthy serum, performed using KOBAS 3.0. Pathways enriched in upregulated genes are shown in red, while pathways enriched in downregulated genes are shown in blue. (B) Bubble plot of GO pathway enrichment analysis of DEGs identified in ECs isolated from SSc BVOs treated with SSc_aDU serum compared to healthy serum, performed using KOBAS 3.0. Pathways enriched in upregulated genes are shown in red, while pathways enriched in downregulated genes are shown in blue. (C-D) TF activity analysis in whole SSc BVO (C) or ECs purified from SSc BVOs (D) exposed to SSc_aDU serum versus healthy serum, with TF with increased activity upon exposure to SSc_aDU serum shown in red, and TF with reduced activity shown in blue. (E) CODEX-based protein expression levels of TWIST1, ZEB1, SNAI1, SNAI2, αSMA, FRA2, ICAM, and COL1 in healthy BVOs treated with healthy serum, SSc_aDU serum or SSc_noDU serum (n≥5 from four iPCS clones), shown as box plots (median ± IQR). Statistical significance was determined by the Kruskal-Wallis test followed by the Dunn multiple comparisons test. SSc: systemic sclerosis; BVO: blood vessel organoids; DAR: differentially accessible region; DEG: differentially expressed gene; SSc_aDU serum: serum from SSc patients with active digital ulcers; TF: transcription factor; CODEX: co-detection by indexing; IQR: interquartile range.

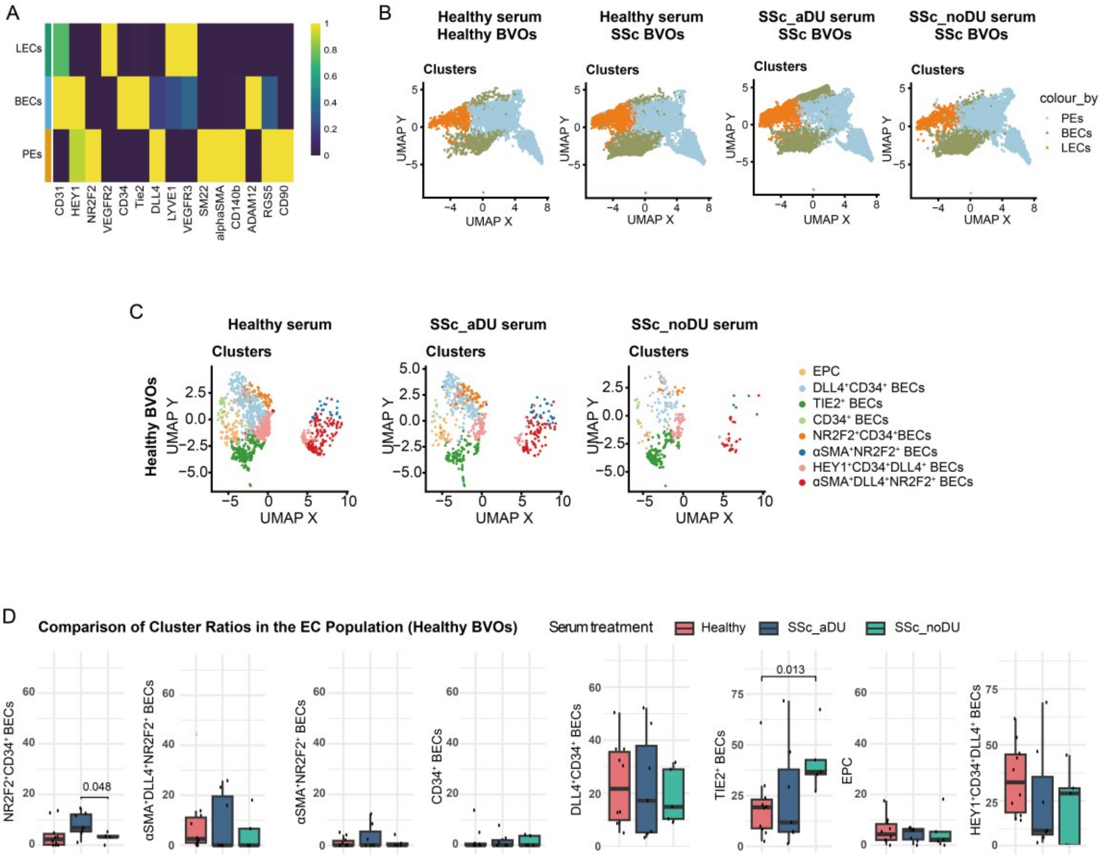


**Figure S4. Main populations in BVOs and shifts in endothelial cell subsets in healthy BVOs upon treatment with SSc serum.** (A) Heatmap showing the relative expression levels of markers for phenotyping of endothelial cells and pericytes across the main populations of LECs, BECs and PEs in BVOs identified by CODEX analysis. (B) UMAP plots showing the BEC subpopulations in healthy BVOs treated with healthy serum, SSc_aDU serum, or SSc_noDU serum. (D) Changes in frequencies of BEC subpopulations in SSc BVOs exposed to healthy serum, SSc_aDU serum or SSc_noDU serum (n≥5 from four iPCS clones) shown as box plots (median ± IQR). Statistically significant p-values (Kruskal-Wallis test followed by Dunn's multiple comparisons test) are included. SSc: systemic sclerosis; BVO: blood vessel organoids; LEC: lymphatic endothelial cells; BEC: blood endothelial cells; PE: pericytes; CODEX: co-detection by indexing; SSc_aDU serum: serum from SSc patients with active digital ulcers; SSc_noDU serum: serum from SSc patients without active digital ulcers; UMAP: Uniform Manifold Approximation and Projection; IQR: interquartile range.

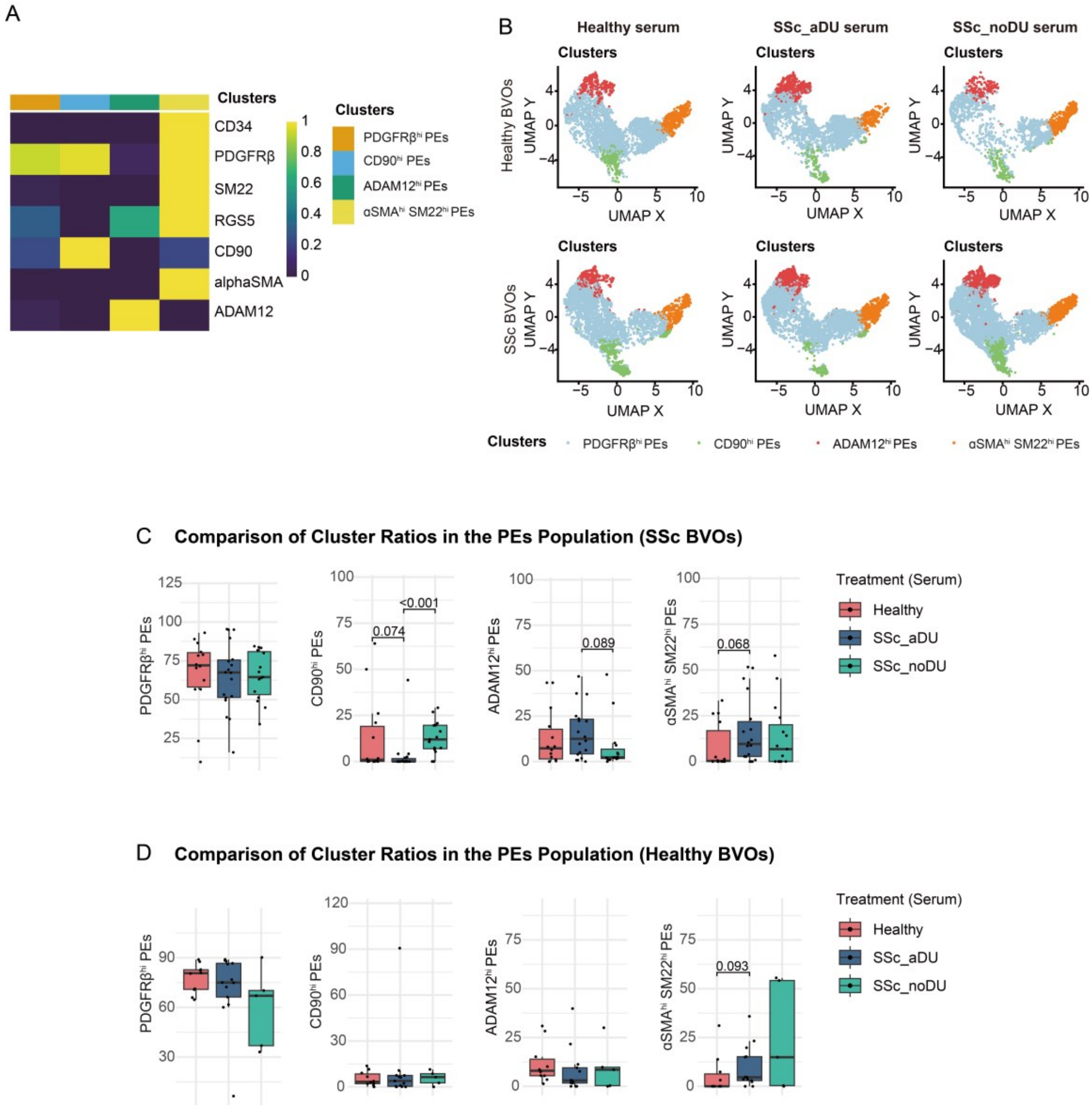


**Figure S5. CODEX-based characterization of pericyte subpopulations in SSc BVOs and their shifts upon treatment with SSc serum.** (A) Heatmap showing the relative expression levels of PE phenotype markers identified by CODEX analysis. (B) UMAP plots showing the PE subpopulations in healthy BVOs (upper row) and SSc BVOs (lower row) treated with healthy serum, SSc_aDU serum or SSc_noDU serum. (C-D) Changes in frequencies of PE subpopulations in SSc BVOs (C) or healthy BVOs (D) exposed to healthy serum, SSc_aDU serum or SSc_noDU serum (n≥11 for SSc BVOs and n≥5 for healthy BVOs from four SSc and four healthy iPCS clones) shown as box plots (median ± IQR). Statistically significant p-values (Kruskal-Wallis test followed by Dunn's multiple comparisons test) are included. SSc: systemic sclerosis; BVO: blood vessel organoids; PE: pericytes; CODEX: co-detection by indexing; SSc_aDU serum: serum from SSc patients with active digital ulcers; SSc_noDU serum: serum from SSc patients without active digital ulcers; UMAP: Uniform Manifold Approximation and Projection; IQR: interquartile range.

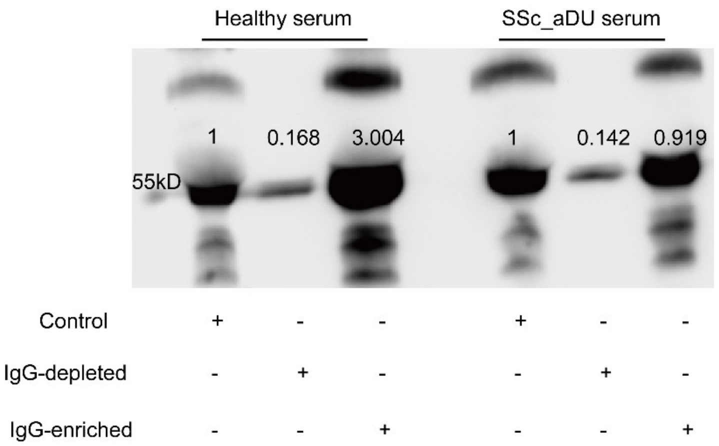


**Figure S6. Efficiency of IgG depletion from human serum.** Western Blot illustrating the presence of IgG protein (55 kD) under different experimental conditions: control (complete serum), IgG-depleted serum, and IgG-enriched serum (IgG concentrated and retained in an agarose-based gel), for both healthy and SSc_aDU serum. Relative band intensities are normalized to the control.

**Table S1. Demographics of iPSC donors**

| Donor | Group | Age | Gender |
|---|---|---|---|
| 1 | healthy | 66 | Female |
| 2 | healthy | 54 | Female |
| 3 | SSc | 47 | Male |
| 4 | SSc | 66 | Male |

**Table S2. CODEX Antibody Panel**

| Target | Clone | Manufacturer | Catalog | Fluoro-phore | Dilution (1:x) |
|---|---|---|---|---|---|
| CD31 | Polyclonal | R&D | AF3628 | ATTO550 | 1500 |
| CD34 | QBEnd/10+ HPCA1/763 | Novus | NBP2-47909 | ATTO647 | 400 |
| αSMA | 1A4 | Biolegend | 614852 | Cy7 | 30000 |
| Podoplanin | D2-40 | Biolegend | 916606 | ATTO550 | 500 |
| CD45 | 2B11 + PD7/26 | Novus | NBP2-34528 | ATTO647 | 900 |
| vWF (1) | Polyclonal | Millipore | AB7356-25UG | Cy7 | 1000 |
| HEY1 | Polyclonal | Thermo | PA5-40553 | ATTO550 | 50 |
| CD90 | EPR3132 | Abcam | ab181885 | ATTO647 | 800 |
| ERG | Polyclonal | Novus | NBP2-60655 | Cy7 | 50 |
| LYVE1 | Polyclonal | R&D | AF2089 | ATTO550 | 100 |
| SNAI1 | Polyclonal | Novus | NBP2-27293 | ATTO647 | 600 |
| SM22 | Polyclonal | Abcam | ab14106 | Cy7 | 3000 |
| ADAM12 | Polyclonal | PROTEINTECH | 14139-1-AP | ATTO550 | 100 |
| VEGFR2 | 55B11 | CST | 28521SF | ATTO647 | 2000 |
| ZEB1 | 3G6 | Invitrogen | 14-9741-80 | Cy7 | 100 |
| CD133 | AC133 | Miltenyi | 130-090-422 | ATTO550 | 50 |
| VEGFR3 | Polyclonal | R&D | AF349 | ATTO647 | 200 |
| VE-cadherin | 123413 | R&D | MAB9381 | Cy7 | 50 |
| PDGFRb | 28E1 | CST | 45176SF | ATTO550 | 200 |
| Cytokeratin | C11 | Biolegend | 628602 | ATTO647 | 4800 |
| Fli-1 | G146-222 (RUO) | BD biosciences | 554266 | Cy7 | 75 |
| FRA2 | 2B2 | Thermo | H00002355-M03 | ATTO550 | 50 |
| Cleaved Caspase 3 | 5A1E | CST | 94530SF | ATTO647 | 200 |
| PROX1 | EPR19273 | Abcam | ab236026 | Cy7 | 50 |
| NR2F2 | Polyclonal | Thermo | PA5-87748 | ATTO550 | 75 |
| Collagen I | EPR7785 | Abcam | ab138492 | ATTO647 | 4500 |
| Tie-2 | Polyclonal | R&D | AF313 | ATTO550 | 150 |
| ICAM | 1A29 | Novus | NBP2-22541-0.1mg | ATTO647 | 50 |
| VCAM | 1.4C3 | Novus | NBP2-33182 | ATTO550 | 150 |
| NG2 | 9.2,27 | Thermo | 14-6504-82 | ATTO647 | 25 |
| SNAI2 | OTI1A6 | Origene | CF800167 | ATTO550 | 150 |
| RGS5 | Polyclonal | PROTEINTECH | 11590-1-AP | ATTO647 | 200 |
| Collagen IV | EPR20966 | Abcam | ab226485 | ATTO550 | 600 |
| Twist-1 | Polyclonal | R&D | AF6230 | ATTO647 | 1000 |
| DLL4 | Polyclonal | PROTEINTECH | 21584-1-AP | ATTO647 | 150 |
